# RavA-ViaA links *Vibrio cholerae* Cpx- and Zra2- envelope stress to antibiotic response

**DOI:** 10.1101/2023.04.24.538083

**Authors:** Evelyne Krin, André Carvalho, Manon Lang, Anamaria Babosan, Didier Mazel, Zeynep Baharoglu

## Abstract

RavA-ViaA were reported to play a role in aminoglycoside sensitivity but the mechanisms remain elusive. Here, we performed competition and survival experiments to confirm that deletion of *ravA-viaA* increases tolerance of the Gram-negative pathogen *Vibrio cholerae* to low and high aminoglycoside concentrations, during aerobic growth. Using high throughput strategies in this species, we identify Cpx and Zra2 two-component systems as new partners of RavA-ViaA. We show that the aminoglycoside tolerance of *Δravvia* requires the presence of these membrane stress sensing two-component systems. We propose that deletion of the RavA-ViaA function facilitates the response aminoglycosides because of a pre-activated state of Cpx and Zra2 membrane stress response systems. We also find an impact of these genes on polymyxin B sensitivity and vancomycin resistance, and we show that simultaneous inactivation of *ravvia* function together with envelope stress response systems leads to outer membrane permeabilization. Vancomycin is mostly used for Gram-positive because of its low efficiency for crossing the Gram-negative outer membrane. Targeting of the *ravA-viaA* operon for inactivation could be a future strategy to allow uptake of vancomycin into multidrug resistant Gram-negative bacteria.

## Introduction

Antibiotic resistance is a growing public health problem. Common resistance mechanisms developed by bacteria consist of limiting antibiotic entry, increasing efflux, degrading the antibiotic or mutating the target. Decreasing intracellular concentrations of antibiotics is one of the most frequent resistance strategies. The aminoglycoside (AG) class of antibiotics targets the ribosome, leading to mistranslation, protein misfolding and eventually cell death. AGs are highly efficient against Gram-negative bacteria [1]. They comprise kanamycin, tobramycin, gentamicin, neomycin, amikacin and streptomycin and are commonly used worldwide. The mechanism of entry for AG into bacterial cells has been in discussion for many years. The currently accepted model for AG entry [2, 3] starts with the PMF-dependent [4-6] and respiration-dependent [7-9] entry of a small AG quantity, leading to mistranslation by the ribosome. Incorporation of mistranslated proteins would then lead to membrane damage and a subsequent second step AG uptake in large amounts [6, 10].

The *ravA-viaA* operon (formerly *yieMN*) was found to sensitize Gram-negative bacteria to the AGs [11], but the mechanism remains elusive. The presence of RavA-ViaA has been reported mostly in ɣ-proteobacteria. Notably, *ravA-viaA* were found in 37 out of 50 randomly chosen enterobacteria [12]. Structural work revealed that RavA and ViaA interact with each other [13], and form a complex with a third partner, LdcI (or CadA) [12, 13]. It was also shown that the RavA-ViaA complex interacts with phospholipids at the inner membrane [14].

The involvement of *ravA-viaA* genes in AG susceptibility was originally identified by independent approaches in different proteobacteria*, Escherichia coli* [11] and *Vibrio cholerae* [15]. In *E. coli,* overexpression of *ravA-viaA* sensitizes to gentamicin, and its deletion was shown to increase AG resistance [11, 16] but only in anaerobic conditions and low energy state [14, 17]. In *V. cholerae*, the *ravA-viaA* operon (VCA0762-VCA0763) is also involved in the response to low doses (below MIC, or sub-MIC) of AGs, this time in aerobic conditions. While studying the response of *V. cholerae* to sub-MIC tobramycin we have conducted a high throughput transposon insertion sequencing (TN-seq) screen [18, 19] where the most enriched insertions were detected in the VC_A0762 (*viaA*) and VC_A0763 (*ravA*) genes (respectively 60x and 30x enrichment), suggesting that inactivation of the operon confers a growth advantage in the presence of AGs in *V. cholerae.* This is consistent with our previous results showing that genotoxic stress induced by tobramycin is limited in the absence of this operon [15].

The PMF is involved in the first phase of AG uptake [20], while the second phase occurs in response to mistranslation (or through sugar transporters). PMF is produced by the activity of electron transfer chains in respiratory complexes, where Fe-S cluster are key actors. It was shown that Fe-S biogenesis and their fueling to respiratory complexes have a direct impact on AG uptake [21]. Previous studies reported that RavA and ViaA interact with Fe-S cluster biogenesis machineries and also with components of the major Fe-S cluster containing Nuo respiratory complex [11, 22], suggesting that RavA-ViaA could contribute to folding of the respiratory complex I. Therefore, it was proposed that RavA and ViaA sensitize *E. coli* to AGs by facilitating Fe-S targeting to complex I and as a consequence an increased level of PMF. Note that *E. coli* and *V. cholerae* are very dissimilar in respect to the respiration complexes, Fe-S biogenesis and oxidative stress response pathways. For instance, *V. cholerae* lacks the above mentioned Cyo/Nuo complex I, and lacks also the SUF Fe-S biogenesis system used under oxidative stress. RavA-ViaA complex, was thus proposed to allows AGs to accumulate inside the cells, presumably by enhancing their uptake of AGs [17].

In order to shed light into how these genes modulate bacterial susceptibility to AGs, it’s important to understand in which conditions these genes are expressed, which bacterial functions are necessary for AG sensitization, and which bacterial processes are affected by these genes.

Here, we first extensively confirmed the RavA-ViaA dependent AG susceptibility and tolerance phenotypes in *V. cholerae*. Next, high throughput approaches identified the involvement of envelope stress responses in these phenotypes. TN-seq showed that inactivation of *cpxP* repressor, i.e. activation Cpx response, is beneficial in *Δravvia*. Cpx responds to conditions that cause misfolding of inner membrane (IM) and periplasmic proteins, and subsequent membrane defect (for review[23]). Cpx also down-regulates outer membrane proteins (OMPs). In parallel, transcriptomic data in *Δravvia* showed strong induction of VC_1314 and the VC_1315-VC1316 operon which presents similarities with the Cpx and Zra two-component envelope stress response systems. In *E. coli*, ZraSR contributes to antibiotic resistance and is important for membrane integrity [24]. *Δzra* has increased membrane disruption during treatment with membrane targeting antibiotics. Zra chaperone activity is enhanced in the presence of zinc ions. In this case, zinc was proposed to be a marker of envelope stress perturbation where ZraPSR is a sentinel sensing and responding to zinc entry into the periplasm [25].

We find that the AG tolerance conferred by *ravA-viaA* deletion in *V. cholerae* requires the presence of the Cpx system and Zra-like system, which we propose to name *zraP2-zraS2-zraR2.* We further show for the first time that the *ravA-viaA* operon is also involved in the response to polymyxin B and that *ravA-viaA* deletion together with inactivation of Cpx or Zra2 envelope stress responses leads to outer membrane permeabilization and may confer vancomycin sensitivity to Gram-negative bacteria.

## Results

### *ravvia* deletion decreases sub-MIC aminoglycoside susceptibility in *V. cholerae*

In order to evaluate AG related roles of RavA-ViaA, we assessed the effect of *ravA-viaA* deletion (referred to as *Δravvia* below) and overexpression (chromosomal extra-copy, referred to as WT::*ravvia*OE+, where OE stands for overexpression). We first performed competitions against *V. cholerae* WT, in the absence and presence of sub-MIC doses of AGs: tobramycin (TOB) and gentamicin (GEN), and also of antibiotics from families other than AGs: chloramphenicol (CM) that targets translation and ciprofloxacin (CIP) that targets DNA replication.

Competition results show that **(Figure 1A**) (i) *V. cholerae Δravvia* has a growth advantage compared to WT during growth with AGs TOB and GEN (40 and 50% MIC); (ii) *V. cholerae WT::ravviaOE+* has a growth disadvantage compared to WT with TOB and GEN; (iii) no significant effect is observed in the presence of sub-MICs of tested antibiotics other than AGs. This suggests the specificity of the mechanism of action of RavA-ViaA to AGs, in *V. cholerae*. Note that the beneficial effect of *Δravvia* is observed here during aerobic growth, while in *E. coli*, it can only be observed in anaerobic conditions.

**Figure 1:**
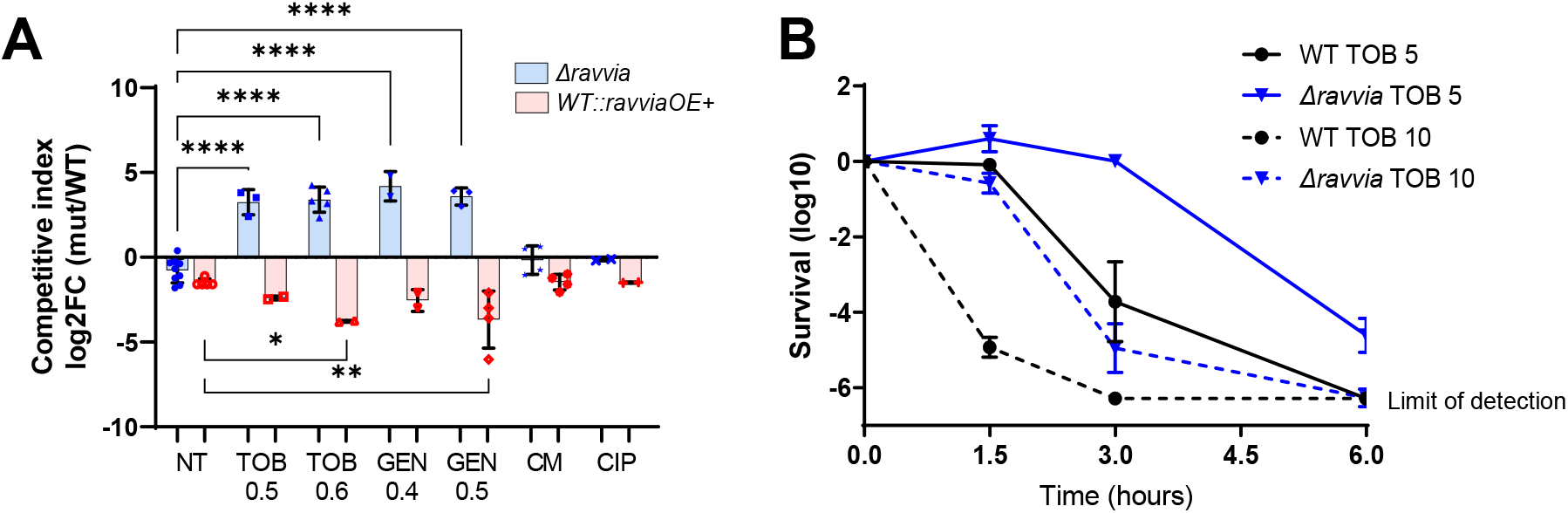
Effect of RavA-ViaA on fitness and tolerance to aminoglycosides. **A.** Competition experiments mixing *ravvia* deletion (Δ) and extra-copy (OE+) mutants and WT in indicated conditions. NT: non-treated. TOB: tobramycin. GEN: gentamicin. CM: chloramphenicol. CIP: ciprofloxacin. The Y-axis represents log_2_ of competitive index value calculated as described in the methods. A competitive index of 1 (i.e. log2 value of 0) indicates equal growth of both strains. For statistical significance calculations, we used one-way ANOVA. **** means p<0.0001, *** means p<0.001, ** means p<0.01, * means p<0.05. Only significant p values are shown. Number of replicates for each experiment: 3<n<8. Concentrations are indicated in µg/ml. **B.** Survival of indicated strain to lethal tobramycin treatment. *V. cholerae* WT and deletion mutant cultures were grown without antibiotics up to early exponential phase, and serial dilutions were plated on MH medium without antibiotics. Exponential phase cultures were then treated with antibiotics at lethal concentrations for the indicated times. At each time point, dilutions were spotted on MH. Y-axis shows survival calculated as number of colonies at time TN divided by the initial number of colonies before antibiotic treatment. TOB: tobramycin 5 or 10 µg/ml.

### *ravvia* deletion increases tolerance to high doses of aminoglycosides in *V. cholerae*

Next, we tested the effect of *ravvia* deletion on resistance by measuring the minimal inhibitory concentration (MIC), and on survival to lethal treatment with AGs. The MIC of *Δravvia* seemed unchanged, or slightly increased compared to WT (1.2 to 1.5 µg/ml instead of 1.2 µg/ml) (**Table 1**). Single Δ*ravA* or Δ*viaA* show the same MIC as the deletion of the whole operon, as expected, since the two proteins form a complex. The *ravviaOE+* mutant shows lower resistance (MIC at 0,75 µg/ml) (**Table 1**).

**Table 1.**
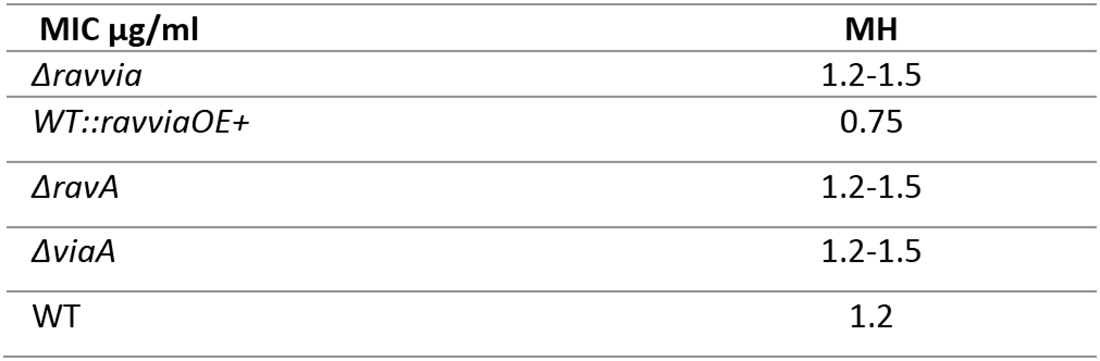

For survival to lethal treatment, we tested the tolerance of the strains to 5x and 10xMIC TOB doses for 6 hours. Deletion of *ravvia* strongly increases survival to lethal TOB doses, when compared to the WT strain (**Figure 1B**). No effect is observed upon treatment with CIP, Trimethoprim and Carbenicillin (**Figure S1).** Treatment to TOB 5 µg/ml during 3 hours shows that *Δravvia* is not affected by TOB during this treatment window (**Figure 1B),** but these cells still die upon longer treatment periods (6 hours) (**Figure 1B)**, excluding any bacteriostatic action of AGs in *Δravvia* mutant. Thus, *Δravvia* is killed upon lethal AG treatment, even though at a much slower rate than the WT, which is the definition of a tolerant population [26].

### The effect of *ravvia* is only partly due to differential aminoglycoside uptake

We tested whether RavA-ViaA complex impacts AG entry, using the AG neomycin coupled to the fluorophore Cy5 (Neo-cy5), as previously done [27, 28]. In this assay, cell fluorescence is proportional to Neo-cy5 uptake. AG uptake increased in *ravvia*OE+ (**Figure 2A**), suggesting that RavA-viaA overexpression facilitates AG entry into the bacterial cell. Surprisingly, AG uptake is not decreased in *V. cholerae Δravvia* (**Figure 2A**). Since AG uptake depends on PMF, we tested the effect of *Δravvia* on PMF. We used Mitotracker assay [28, 29], based on a fluorescent dye which accumulates inside the cell in a PMF dependent way. We observed increased PMF in *ravvia*OE+ strain (**Figure 2B**), strengthening the notion that increased AG entry of *ravvia*OE+ is due to increased PMF. Conversely, no significant decrease of PMF was detected in the AG tolerant Δ*ravvia* strain (**Figure 2B**), suggesting that the effect of *ravvia* on AG susceptibility may require additional explanation as simply PMF modulation. Moreover, as expected, *sdh* (succinate dehydrogenase) deletion increased fitness in AGs, because of a decrease in PMF [5]. Simultaneous deletion of *sdh* and *ravvia* shows an additive fitness advantage (**Figure 2C**), suggesting that the mechanisms of increased fitness of *Δravvia* in sub-MIC TOB is not through a common pathway with the *Δsdh*-dependent PMF decrease.

**Figure 2:**
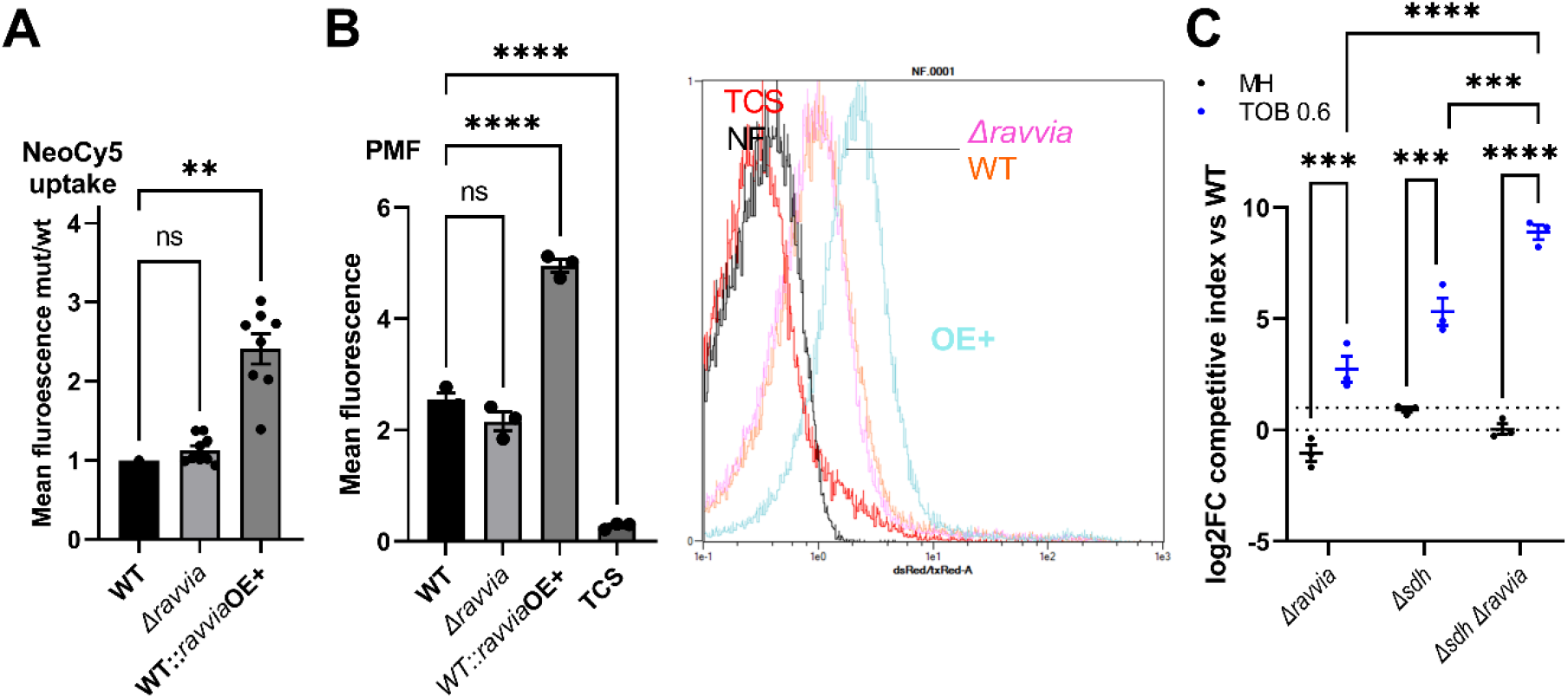
Effect of RavA-ViaA on AG uptake and membrane potential. **A.** Intracellular level of neomycin coupled to the fluorophore Cy5 measured by fluorescence associated flow cytometry. Error bars represent standard deviation. **B.** Quantification of changes in PMF using Mitotracker Red fluorescence measured by flow cytometry. Representative acquisitions are shown: fluorescence is represented in the x-axis (FITC channel), the y-axis represents the number of events corresponding to the number of cells, normalized to height (same number of total cells for both conditions). Each plot represents one experiment. **C.** Competition experiments of *V. cholerae* WT and indicated mutants. MH: no antibiotic treatment (black). TOB: tobramycin 0,6 µg/ml (blue). The Y-axis represents log_2_ of competitive index value calculated as described in the methods. A competitive index of 1 (i.e. log2 value of 0) indicates equal growth of both strains. For statistical significance calculations, we used one-way ANOVA. **** means p<0.0001, *** means p<0.001, ** means p<0.01, * means p<0.05. ns: non-significant. Number of replicates for each experiment: n=3. Only significant p values are shown.

### High throughput approaches point to a role of membrane stress two-component systems in *Δravvia*

In order to further understand changes due to *ravvia* deletion and to search for potential partners of RavA-ViaA, we decided to adopt transcriptomic and TN-seq approaches. RNA-seq was performed on exponentially growing WT and *Δravvia.* Major changes in *Δravvia* compared to WT include more than 10-fold upregulation of sugar transporters, anaerobic respiration, consistent with recent study published by the Barras and Py laboratories [17] and [30], and the VC_1314-1315-1316 genes (**Table 2**).

**Table 2.**
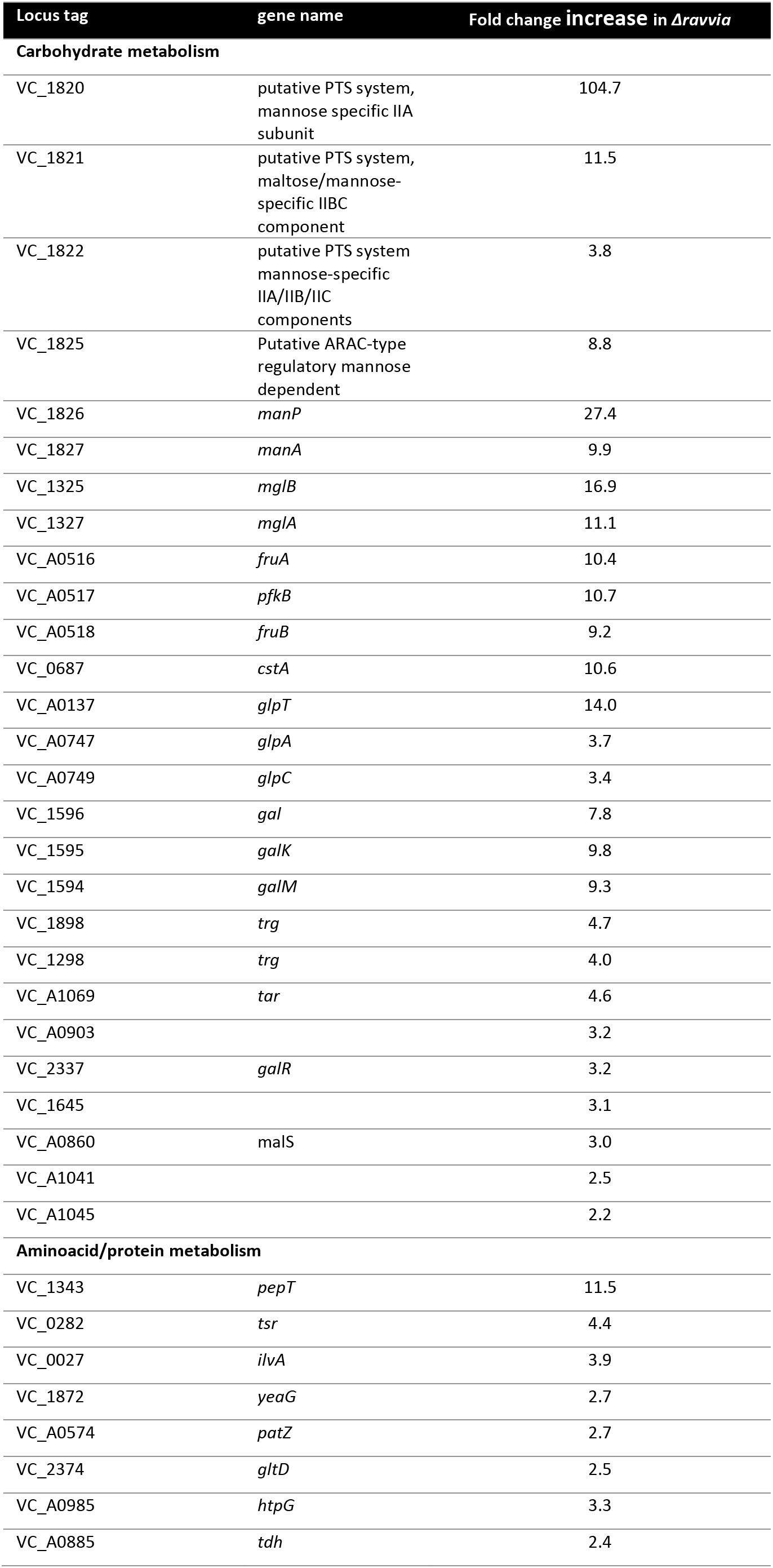
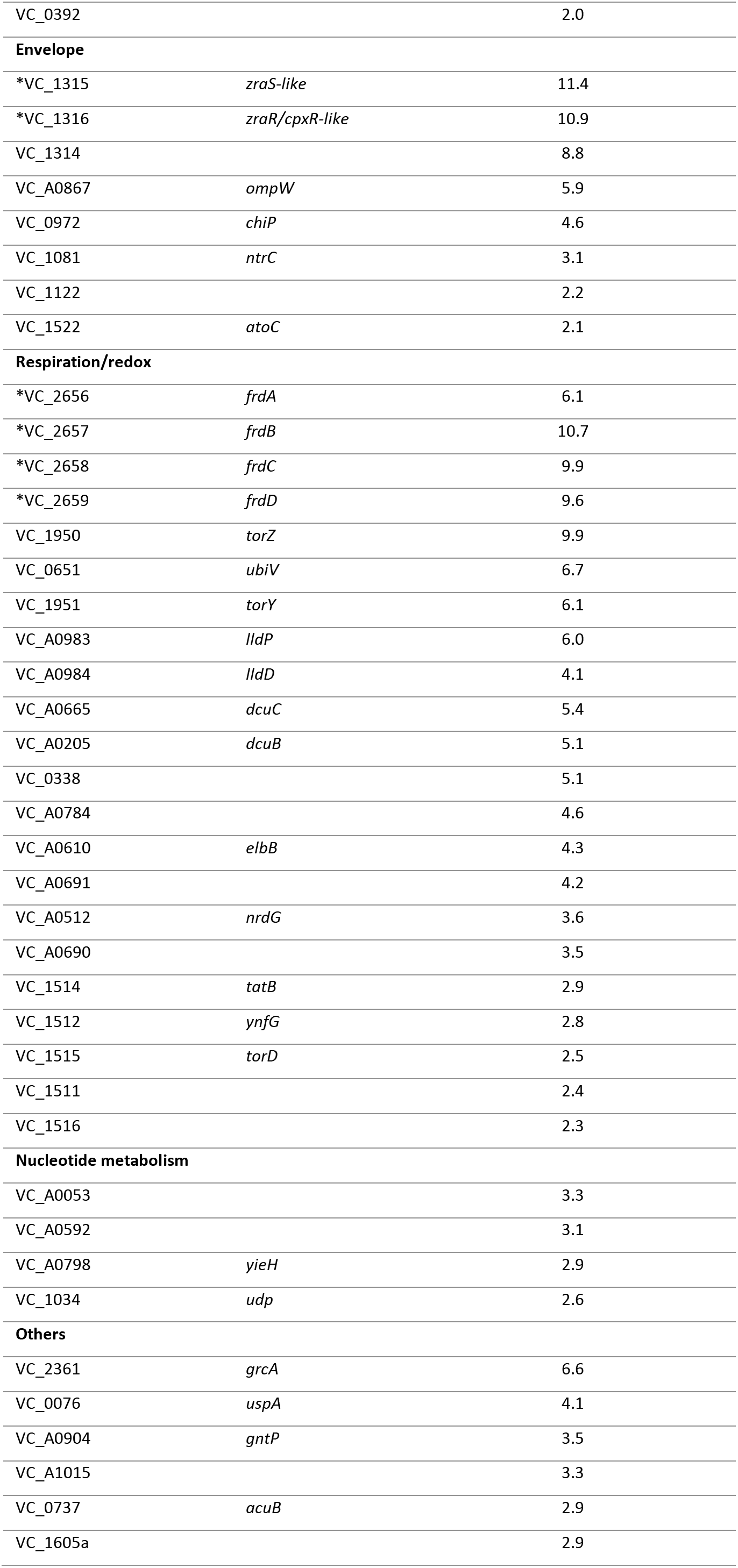
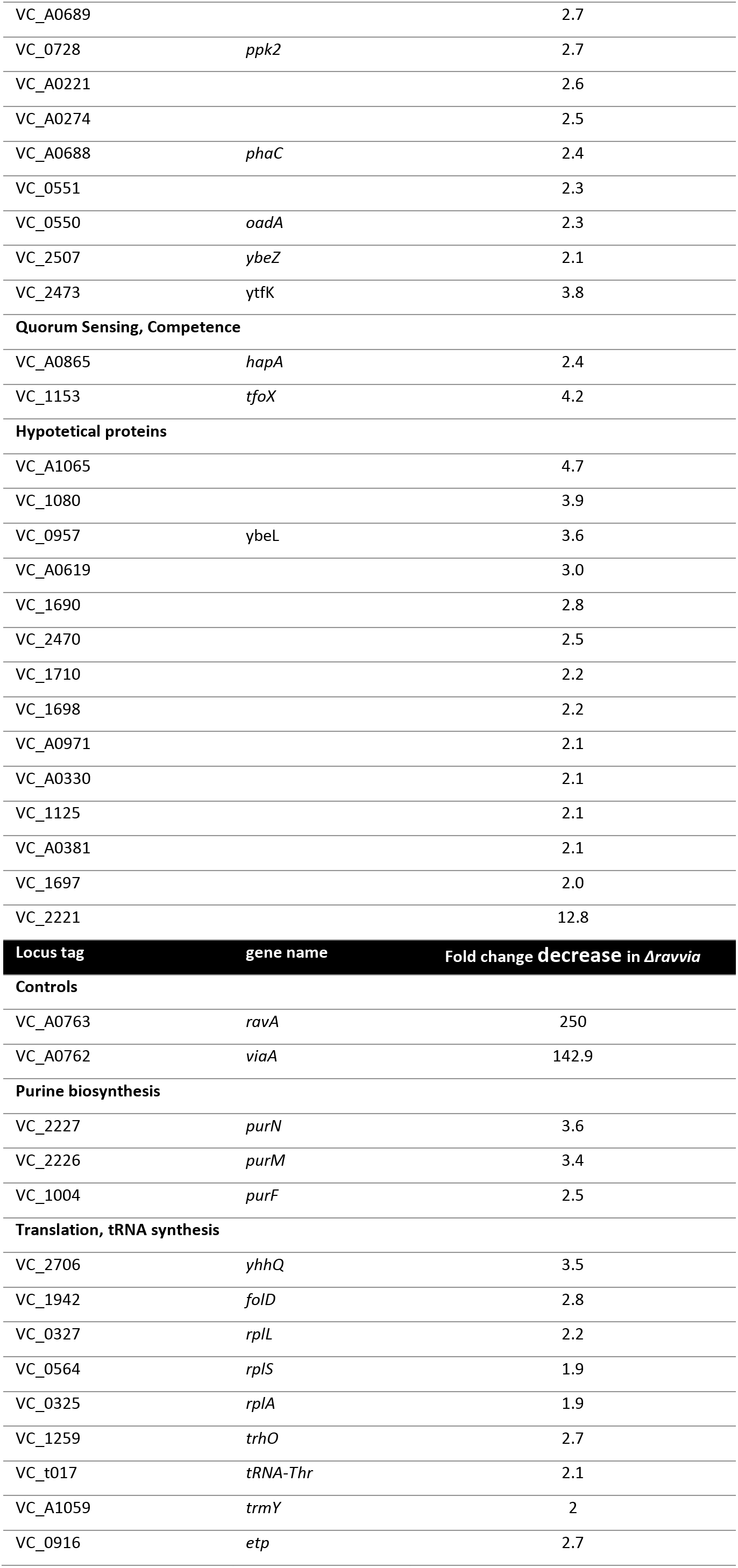
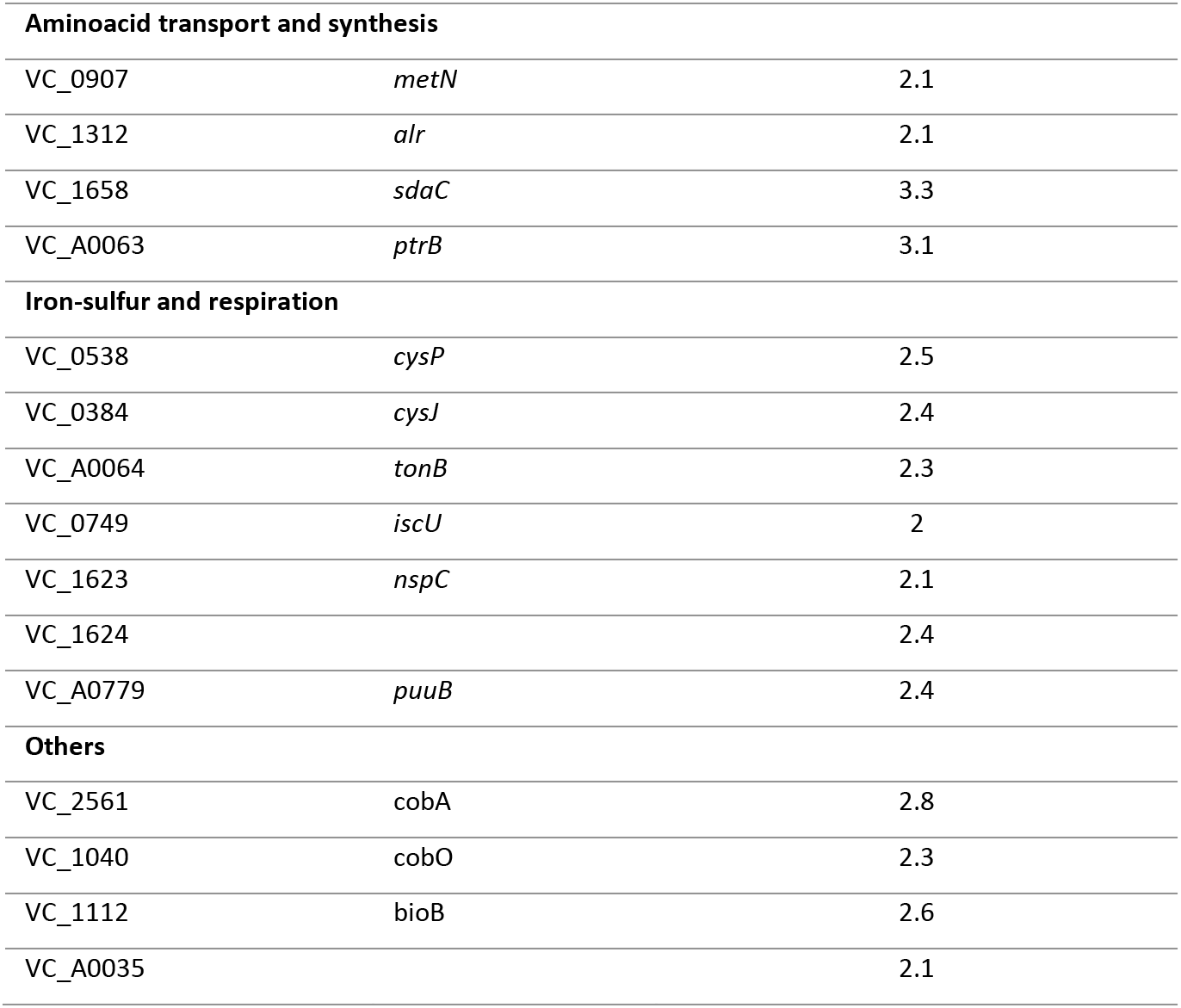
RNA-seq.

The VC_1314 gene and the VC_1315-VC1316 operon show 8- to 10-fold upregulation. VC_1315 presents 35% sequence identity which *E. coli zraS* gene. The zraP regulator and z*raSR* two-component membrane stress response system is involved in antibiotic resistance [24]. VC_1315 also presents 29% (and 21%) sequence identity with the *V. cholerae (*and *E. coli) cpxA* gene. The periplasmic CpxP is the negative regulator of the envelope stress response CpxRA system [31]. VC_1316 presents 26% sequence identity with the *E. coli cpxR* and 23% with the *V. cholerae cpxR*. VC_1314 does not show any sequence similarity neither to *zraP*, nor to *cpxP*. In E. coli, The CpxP-CpxAR system and the ZraP-ZraRS systems, were proposed to be functional homologues [25]. We called VC_1314-1315-1316, the *zra-like* system below in this manuscript.

In parallel to the transcriptomic study, we applied our previously described comparative TN-seq approach [15, 18], to *V. cholerae Δravvia,* to search for genes that are important for survival in the presence of sub-MIC TOB, again in aerobic conditions. We sequenced mutant libraries before and after 16 generations without and with TOB at 50% of the MIC. After sequencing, comparative analysis of the number of detected gene inactivations between the two conditions, indicates whether a given gene is important for growth in the antibiotic (decreased number of reads), or wether its inactivation is beneficial (increased number of raeds), or unchanged. **Tables 3, 4 and 5** show the exhaustive lists of at least 2-fold differentially detected genes with transposon insertions in WT and *Δravvia*, with and without TOB. Deletion of *ravvia* leads to changes in factors involved in carbon metabolism, iron and respiration, and membrane stress. We constructed deletion mutants in WT and Δ*ravvia* contexts and performed competition experiments **(Figure S2**) for 21 of these genes to validate TN-seq results and to identify factors necessary for AG tolerance of *Δravvia*. Competition results were mostly consistent with TN-seq data. Among identified factors, one, *cpxP*, has particularly caught our attention because its inactivation is beneficial in *Δravvia* **(Figure S2**), and because inactivation of *ravvia* in *ΔcpxP* does not cause an additional increase in fitness, suggesting that they may act in the same pathway. Because these systems were also identified as induced in the *ravvia* transcriptome analysis above, we decided to focus on the link between the AG tolerant phenotypes of *Δravvia* and envelope stress response through Cpx and Zra-like systems.

**Table 3.**
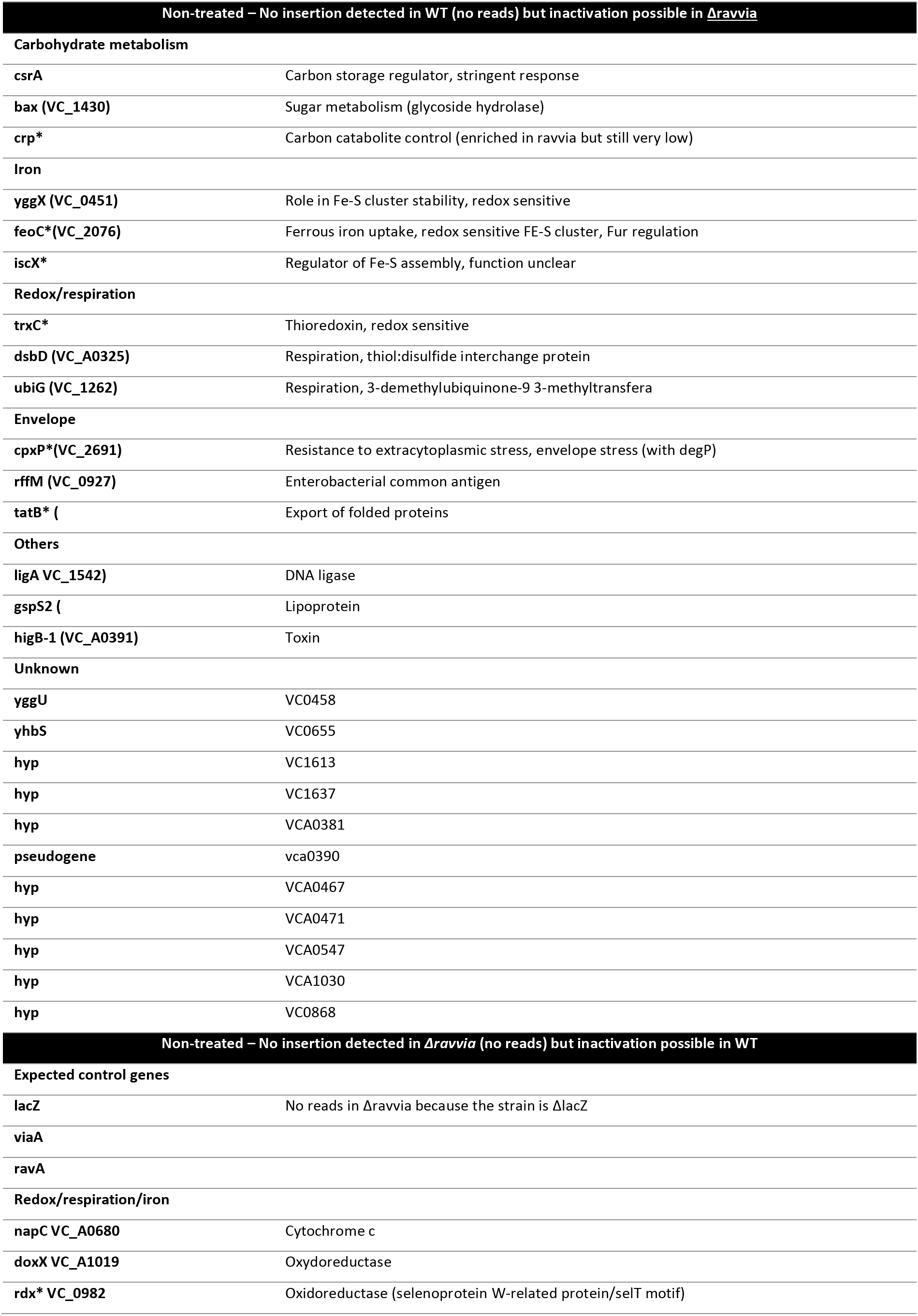
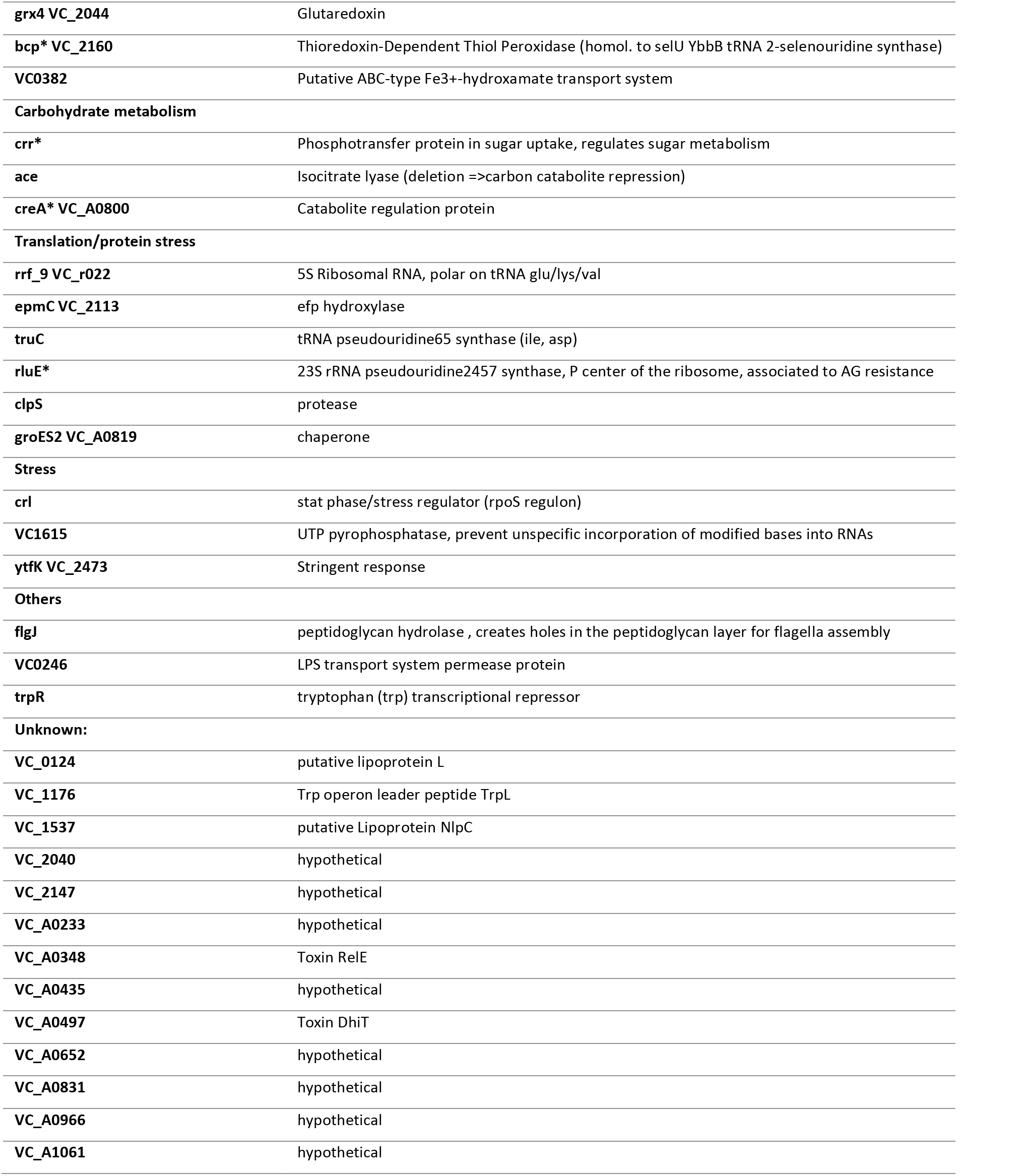
Transposon insertion sequencing at time T0: genes that cannot be inactivated only in WT or only in *Δravvia*.

**Table 4.** Transposon insertion sequencing at time T16, with no treatment: genes that tolerate more insertions either in WT or in *Δravvia*.

| Non- treated 16 gen–<br>More insertions in ravvia |  |  |  |  |  |  |  |  |  |  |
| --- | --- | --- | --- | --- | --- | --- | --- | --- | --- | --- |
| gene name |  | T0<br>ravvia | T16<br>ravvia | T16 TOB<br>ravvia | T0 WT | T16 WT | T16 TOB<br>WT | FC T0<br>ravvia/wt | FC T16<br>ravvia/wt | FC T16<br>TOB<br>ravvia/wt |
| carbohydrate metabolism |  |  |  |  |  |  |  |  |  |  |
| araD | VC_A0244 | 194 | 174 | 143 | 58 | 110 | 97 | 3.3 | 1.6 | 1.5 |
| malK |  | 510 | 242 | 352 | 140 | 96 | 142 | 3.6 | 2.5 | 2.5 |
|  | VC_0549 | 482 | 232 | 334 | 152 | 222 | 173 | 3.2 | 1.0 | 1.9 |
| redox |  |  |  |  |  |  |  |  |  |  |
|  | VC_A0506 | 234 | 88 | 212 | 69 | 165 | 108 | 3.4 | 0.5 | 2.0 |
| nucleotide pool |  |  |  |  |  |  |  |  |  |  |
| surE/umpG |  | 409 | 352 | 323 | 140 | 152 | 69 | 2.9 | 2.3 | 4.7 |
| pnpP | VC_A0970 | 331 | 212 | 204 | 91 | 241 | 132 | 3.6 | 0.9 | 1.5 |
| envelope/membrane |  |  |  |  |  |  |  |  |  |  |
|  | VC_A0152 | 410 | 422 | 344 | 112 | 320 | 150 | 3.7 | 1.3 | 2.3 |
| MshF | VC_0407 | 869 | 525 | 618 | 154 | 127 | 151 | 5.6 | 4.1 | 4.1 |
| VpsQ | VC_0939 | 901 | 679 | 783 | 292 | 313 | 283 | 3.1 | 2.2 | 2.8 |
| others |  |  |  |  |  |  |  |  |  |  |
| mobA |  | 212 | 141 | 151 | 63 | 67 | 71 | 3.4 | 2.1 | 2.1 |
| citX |  | 411 | 331 | 311 | 117 | 203 | 86 | 3.5 | 1.6 | 3.6 |
| trpD |  | 149 | 142 | 187 | 63 | 60 | 59 | 2.4 | 2.3 | 3.2 |
| phhB | VC_A0827 | 546 | 391 | 384 | 154 | 178 | 148 | 3.5 | 2.2 | 2.6 |
| unknown |  |  |  |  |  |  |  |  |  |  |
| alpA | VC_1809 | 286 | 373 | 383 | 90 | 112 | 183 | 3.2 | 3.3 | 2.1 |
| YebG | VC_2326 | 340 | 237 | 286 | 107 | 156 | 110 | 3.2 | 1.5 | 2.6 |
| ynjD | VC_1666 | 206 | 225 | 222 | 32 | 98 | 101 | 6.3 | 2.3 | 2.2 |
| hyp | VC_A0743 | 259 | 241 | 258 | 50 | 81 | 67 | 5.2 | 3.0 | 3.9 |
| hyp | VC_A0440 | 353 | 355 | 421 | 78 | 139 | 69 | 4.5 | 2.6 | 6.1 |
| yaeP | VC_0872 | 395 | 348 | 271 | 91 | 184 | 201 | 4.3 | 1.9 | 1.3 |
| hyp | VC_A0086 | 297 | 271 | 267 | 71 | 254 | 191 | 4.2 | 1.1 | 1.4 |
| hyp | VC_A0649 | 360 | 331 | 318 | 93 | 188 | 195 | 3.9 | 1.8 | 1.6 |
| hyp | VC_A0032 | 516 | 417 | 542 | 137 | 158 | 114 | 3.8 | 2.6 | 4.8 |
| ? | VC_A0429-30 | 195 | 262 | 375 | 52 | 113 | 149 | 3.7 | 2.3 | 2.5 |
| hyp | VC_A1030 | 173 | 137 | 115 | 55 | 40 | 58 | 3.1 | 3.5 | 2.0 |
| YecM | VC_2073 | 191 | 205 | 181 | 61 | 134 | 120 | 3.1 | 1.5 | 1.5 |
| Non treated 16 gen–<br>more insertions in WT |  |  |  |  |  |  |  |  |  |  |
| gene name |  | T0<br>ravvia | T16<br>ravvia | T16 TOB<br>ravvia | T0 WT | T16 WT | T16 TOB<br>WT | FC T0<br>wt/ravvia | FC T16<br>wt/ravvia | FC T16<br>TOB<br>wt/ravvia |
| Translation/protein |  |  |  |  |  |  |  |  |  |  |
| yjgA | VC_2536 | 51 | 96 | 77 | 163 | 172 | 142 | 3.2 | 1.8 | 1.8 |
| membrane transport |  |  |  |  |  |  |  |  |  |  |
|  | VC_1605a | 16 | 31 | 38 | 198 | 146 | 99 | 12.5 | 4.7 | 2.6 |
| pspG |  | 56 | 194 | 113 | 561 | 579 | 421 | 10.0 | 3.0 | 3.7 |
|  |  | 38 | 6 | 25 | 148 | 22 | 27 | 3.9 | 3.4 | 1.1 |
| yhdZ | VC_A1037 | 55 | 93 | 128 | 177 | 165 | 165 | 3.2 | 1.8 | 1.3 |
| carbohydrate metabolism |  |  |  |  |  |  |  |  |  |  |
| pfkA |  | 31 | 13 | 28 | 124 | 54 | 62 | 4.0 | 4.0 | 2.2 |
| galU |  | 68 | 48 | 23 | 215 | 100 | 16 | 3.2 | 2.1 | 0.7 |
| regulators |  |  |  |  |  |  |  |  |  |  |
| CspA | VC_A0166 | 30 | 24 | 28 | 115 | 162 | 32 | 3.8 | 6.7 | 1.1 |
| **marR** | VC_A1005 | 37 | 92 | 65 | 181 | 190 | 110 | 4.8 | **2.1** | **1.7** |
|  | VC_A0999 | 65 | 65 | 69 | 239 | 144 | 123 | 3.7 | **2.2** | **1.8** |
| **FE-S/redox** |  |  |  |  |  |  |  |  |  |  |
| **DsbD** | VC_A0389 | 39 | 99 | 101 | 155 | 195 | 103 | 4.0 | **2.0** | **1.0** |
| **ibaG** | VC_2515 | 8 | 19 | 9 | 114 | 91 | 6 | 14.3 | **4.8** | **0.7** |
| **grxB** |  | 36 | 43 | 47 | 129 | 119 | 68 | 3.6 | **2.7** | **1.5** |
|  | VC_1254 | 61 | 102 | 190 | 216 | 193 | 387 | 3.5 | **1.9** | **2.0** |
| **hutC** |  | 55 | 72 | 88 | 187 | 164 | 127 | 3.4 | **2.3** | **1.4** |
| **speB** |  | 26 | 41 | 19 | 109 | 73 | 35 | 4.2 | **1.8** | **1.9** |
| **thiS** |  | 63 | 38 | 78 | 202 | 157 | 105 | 3.2 | **4.1** | **1.3** |
| **electron transfer** |  |  |  |  |  |  |  |  |  |  |
| **ccmF** | VC_A0368 | 88 | 136 | 103 | 311 | 224 | 201 | 3.5 | **1.6** | **2.0** |
| **ccmE** |  | 61 | 36 | 81 | 190 | 108 | 846 | 3.1 | **3.0** | **10.4** |
| **cobO** |  | 73 | 77 | 82 | 221 | 180 | 165 | 3.0 | **2.3** | **2.0** |
| **DNA repair** |  |  |  |  |  |  |  |  |  |  |
| **xthA** |  | 37 | 38 | 37 | 140 | 158 | 141 | 3.8 | **4.1** | **3.8** |
| **two component system** |  |  |  |  |  |  |  |  |  |  |
| **dpiB citA\*** | VC_0791 | 48 | 67 | 65 | 179 | 161 | 119 | 3.7 | **2.4** | **1.8** |
| **cheY** | VC_A1096 | 61 | 39 | 62 | 207 | 154 | 36 | 3.4 | **4.0** | **0.6** |
| **others** |  |  |  |  |  |  |  |  |  |  |
| **hfq** |  | 33 | 38 | 0 | 117 | 116 | 0 | 3.5 | **3.1** | **no reads** |
| **unknown** |  |  |  |  |  |  |  |  |  |  |
| **hyp** | VC_2367 | 22 | 103 | 85 | 199 | 173 | 160 | 9.0 | **1.7** | **1.9** |
| **yacL** | VC_0605 | 59 | 125 | 131 | 437 | 371 | 224 | 7.5 | **3.0** | **1.7** |
| **hyp** | VC_A0494 | 24 | 35 | 14 | 113 | 141 | 139 | 4.8 | **4.0** | **10.2** |
| **hyp** | VC_A0919 | 60 | 107 | 68 | 284 | 174 | 260 | 4.7 | **1.6** | **3.8** |
| **ybhG** | VC_1659 | 33 | 78 | 73 | 143 | 154 | 104 | 4.3 | **2.0** | **1.4** |
| **hyp** | VC_A0458 | 92 | 127 | 126 | 369 | 275 | 183 | 4.0 | **2.2** | **1.5** |
| **flit** |  | 75 | 390 | 89 | 296 | 869 | 23 | 4.0 | **2.2** | **0.3** |
| **hyp** | VC_2046 | 143 | 202 | 153 | 539 | 562 | 281 | 3.8 | **2.8** | **1.8** |
| **hyp** | VC_A0387 | 145 | 172 | 183 | 545 | 363 | 288 | 3.7 | **2.1** | **1.6** |
| **rsta2** | VC_1463 | 68 | 17 | 0 | 253 | 205 | 138 | 3.7 | **11.7** | **no reads in ravvia** |
| **wbeT rfbT** | VC_0258 | 37 | 46 | 36 | 136 | 154 | 93 | 3.7 | **3.3** | **2.5** |
| **unfA** | VC_1191 | 48 | 81 | 125 | 176 | 128 | 78 | 3.6 | **1.6** | **0.6** |
| **hyp** | VC_A1024 | 136 | 227 | 217 | 483 | 597 | 369 | 3.6 | **2.6** | **1.7** |
| **hyp** | VC_0491 | 39 | 81 | 68 | 138 | 123 | 90 | 3.5 | **1.5** | **1.3** |
| **hyp** | VC_A0342 | 59 | 112 | 125 | 189 | 133 | 113 | 3.2 | **1.2** | **0.9** |

**Table 5.**
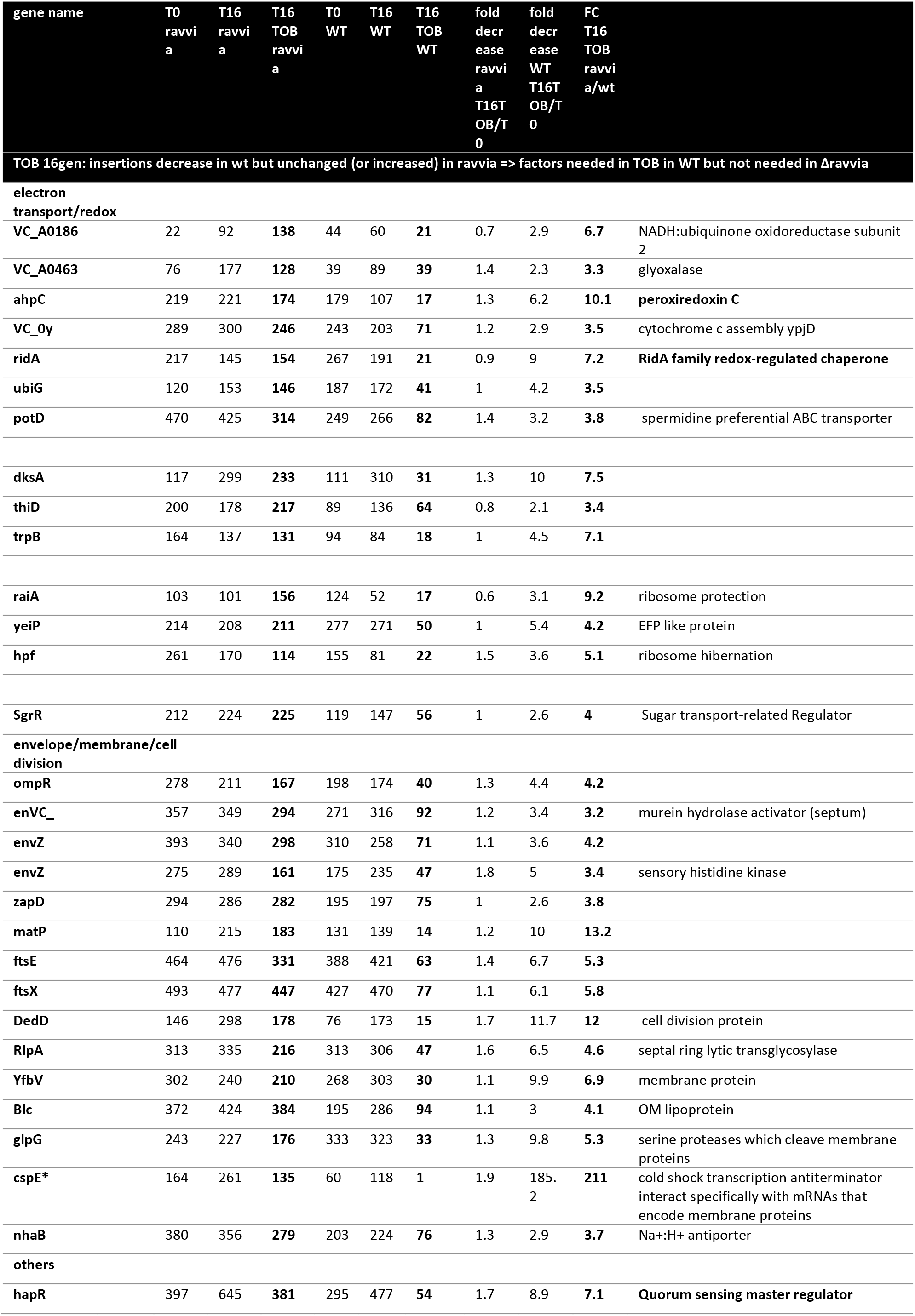
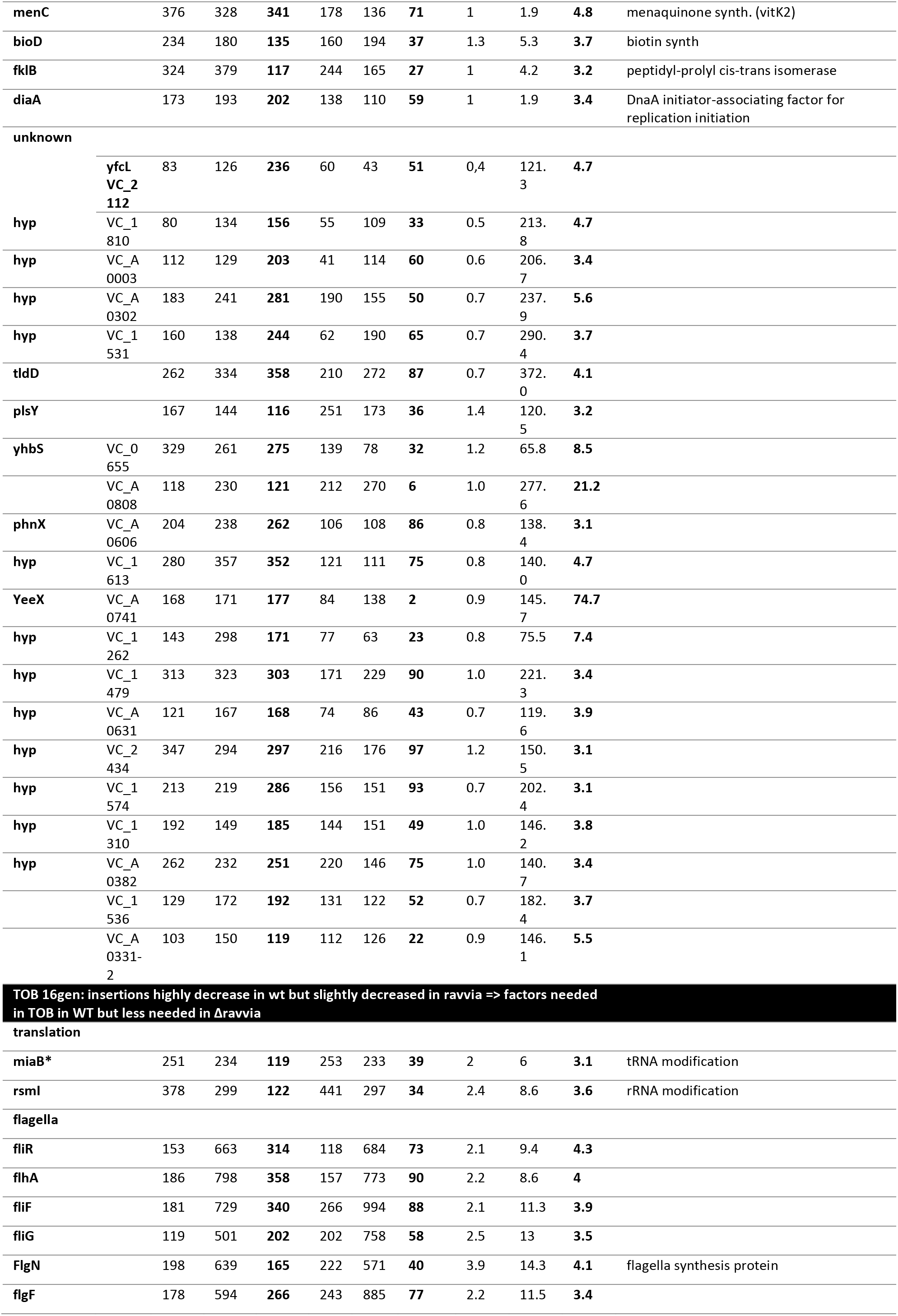
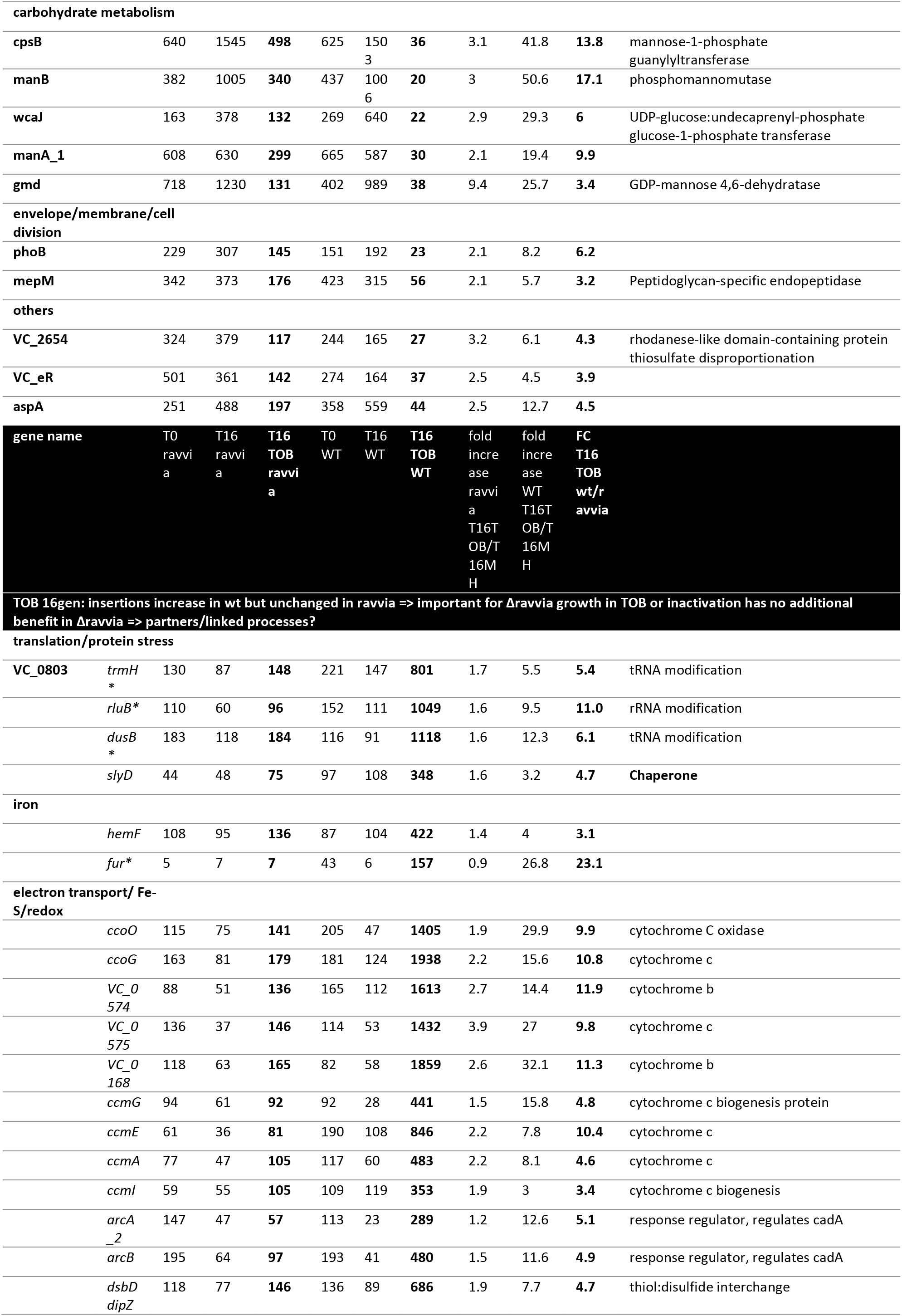
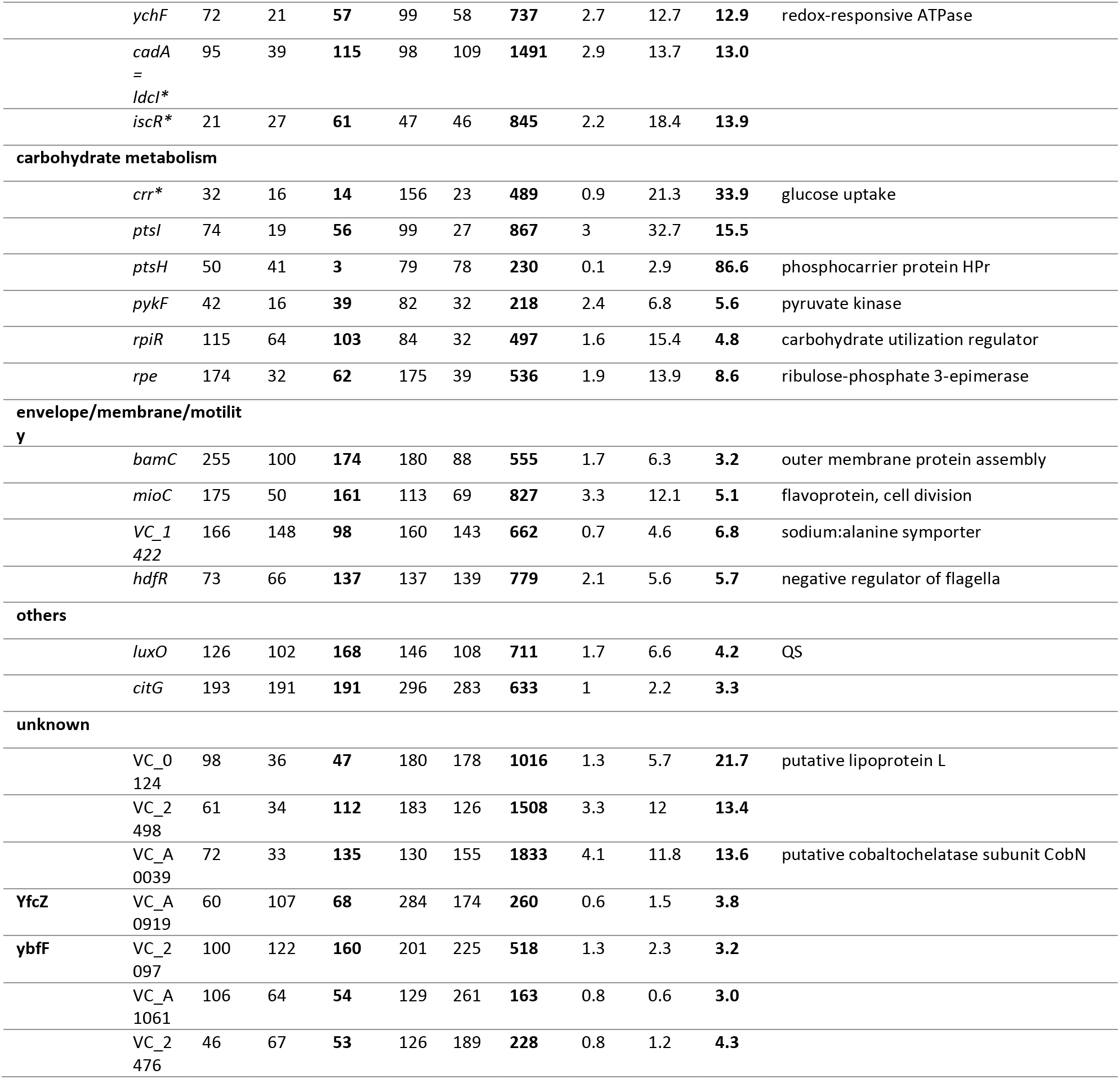
Transposon insertion sequencing at time T16, with sub-MIC TOB: genes that tolerate more insertions either in WT or in *Δravvia*.

### Cpx and Zra-like system mediated envelope stress response systems are necessary for the fitness advantage of *Δravvia* during growth with sub-MIC AGs and AG tolerance

In order to assess the importance of the Cpx and the putative Zra-like systems in the response to AGs of *Δravvia,* we tested fitness and tolerance in competition and survival experiments in the absence of one or both of these systems. **Figure 3ABC** shows competition experiments with *cpx* and *zra-like* operons inactivation. Inactivation of *cpx* alone does not affect fitness in TOB (**Figure 3A**) while inactivation of *zra-like* alone increases fitness in TOB (**Figure 3B**). The fitness advantage of *Δzra-like* depends on the presence of *cpx*, since deletion of *cpx* in *Δzra-like* suppresses its fitness advantage (**Figure 3C**). Strikingly, the deletion of *cpx* or *zra-like* in *Δravvia* leads to, respectively, loss or strong decrease of the fitness advantage of *Δravvia* in TOB (**Figure 3A and B**). The triple mutant *Δravvia Δcpx Δzra* shows a phenotype similar to the *Δravvia Δcpx* double mutant (**Figure 3C**). These results show that Cpx envelope stress response system is necessary for the enhanced tolerance of *Δravvia* to TOB, and that the Zra-like system also contributes significantly to this fitness advantage.

**Figure 3:**
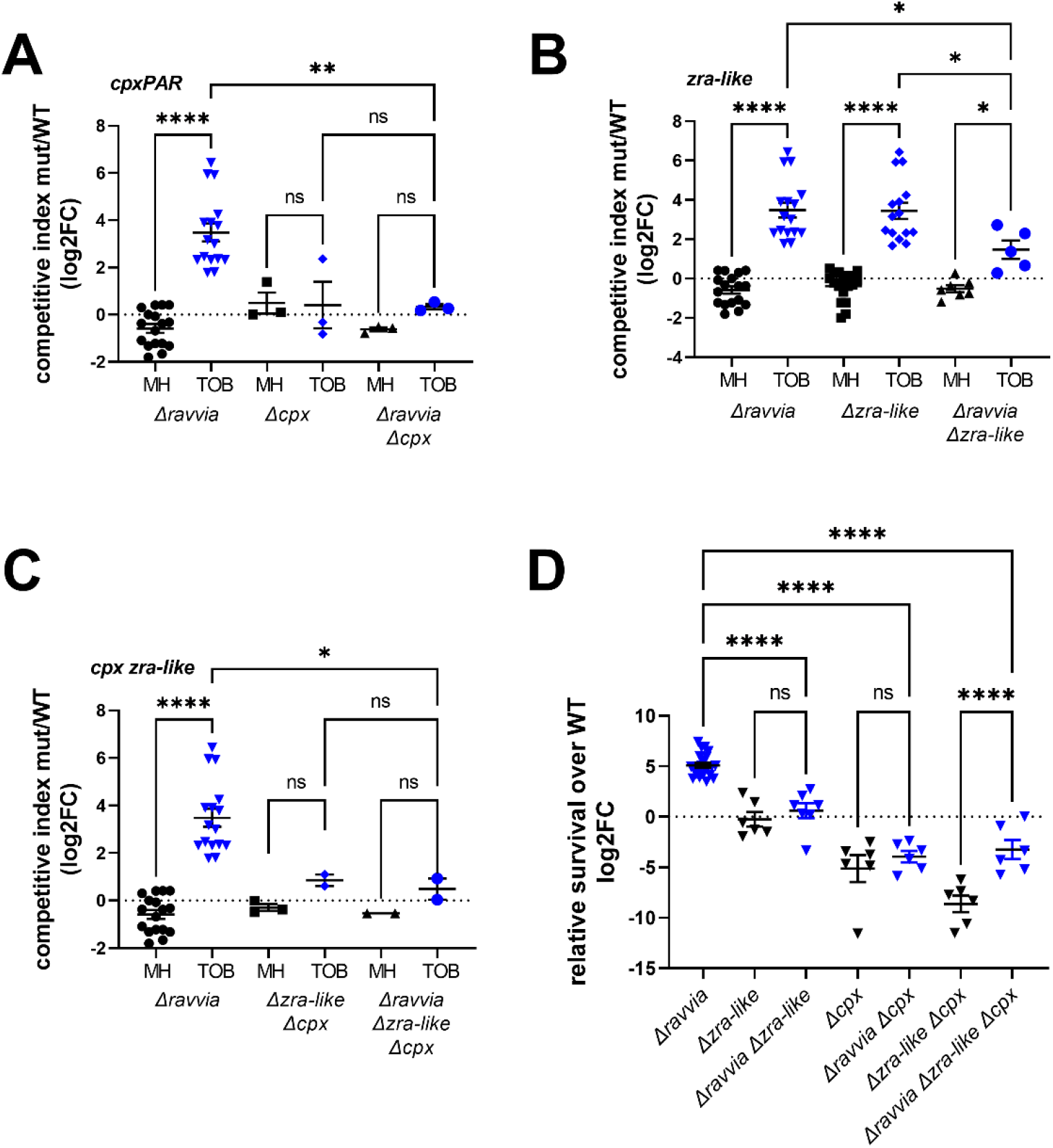
Cpx and Zra-like two component envelope stress response systems are involved in fitness increase of *Δravvia* with TOB and TOB tolerance. ABC. Competitions. The effect of deletion of *cpx* (**A**), or *zra-like* (**B**), or both (**C**) on competitive index in MH without and with TOB, where MH is the untreated growth medium. *In vitro* competition experiments of *V. cholerae* WT and indicated mutants in specified media: in black: MH: no antibiotic treatment. In blue: TOB: tobramycin 0,6 µg/ml. The Y-axis represents log_2_ of competitive index value calculated as described in the methods. A competitive index of 1 (i.e. log2 value of 0) indicates equal growth of both strains. **D. Tolerance.** Cultures were grown to exponential phase in MH medium. Survival of WT and *Δravvia* to 3-hours treatment with lethal TOB at 5x MIC 5 µg/ml was measured. The Y-axis represents log2 value of survival ratios, calculated as survival of the mutant over survival of the WT. A relative survival ratio of 1 (i.e. log2 = 0) indicates equal survival as the WT strain. For statistical significance calculations, we used one-way ANOVA. **** means p<0.0001, *** means p<0.001, ** means p<0.01, * means p<0.05. ns: non-significant. Number of replicates for each experiment: 3<n<8.

We next performed TOB tolerance tests using a concentration of 5x MIC for 3 hours **(Figure 3D)**. Under these conditions, the survival of the single Δ*zra-like* system mutant is slightly lower than WT, the survival of *Δcpx* is lower and both systems seem to be additive as the double mutant appears to show even lower tolerance than the single *Δcpx.* Strikingly, the high level of tolerance of *Δravvia* is completely lost upon deletion of *zra-like* and goes even lower than WT upon deletion of *cpx* and in the triple mutant. For unknown reasons, the decrease of tolerance is stronger in *Δzra-like Δcpx* than in *Δzra-like Δcpx Δravvia*, as if deletion of *zra-like* in *Δravvia Δcpx* was beneficial. In any case, the AG tolerance conferred by the deletion of *ravvia* necessitates the presence of both Cpx and Zra-like systems.

### *Δavvia* related phenotypes are linked to extracellular zinc concentrations

Since Cpx and Zra systems have previously been associated with metals such as iron and zinc, we next performed competition experiments in the presence of these metals. The presence of supplemented iron did not affect the fitness of the *Δravvia* derivatives in any condition (**Figure S3ABC**). Zinc supplementation (**Figure 4ABC**) restores fitness in TOB for the *Δravvia Δcpx* and *Δravvia Δcpx* double mutants but not of the triple mutant, suggesting that the effect of zinc in *Δravvia* is somehow linked to the Cpx and Zra-like systems, which may act in a redundant way.

Moreover, while zinc has no effect on the TOB tolerance phenotype of *Δravvia* or *Δravvia Δzra-like*, it restores high tolerance to the *Δravvia Δcpx* mutant (**Figure 4D**), which is consistent with the zinc-dependent increase of fitness of the *Δravvia Δcpx* mutant shown in **Figure 4C**. We wondered whether these genes could be regulated by zinc. We found that *cpx*, but not *zra*, mRNA levels are increased in the presence of zinc (**Figure 4E**). Ravvia expression from P*ravvia* promoter fused to *gfp* is also induced by zinc (**Figure 4F**). Overall, results indicate that the *Δravvia* mutant’s AG tolerance is dependent on the Cpx and Zra-like systems, and that *ravvia* function is also somehow associated with the sensing of zinc levels.

**Figure 4:**
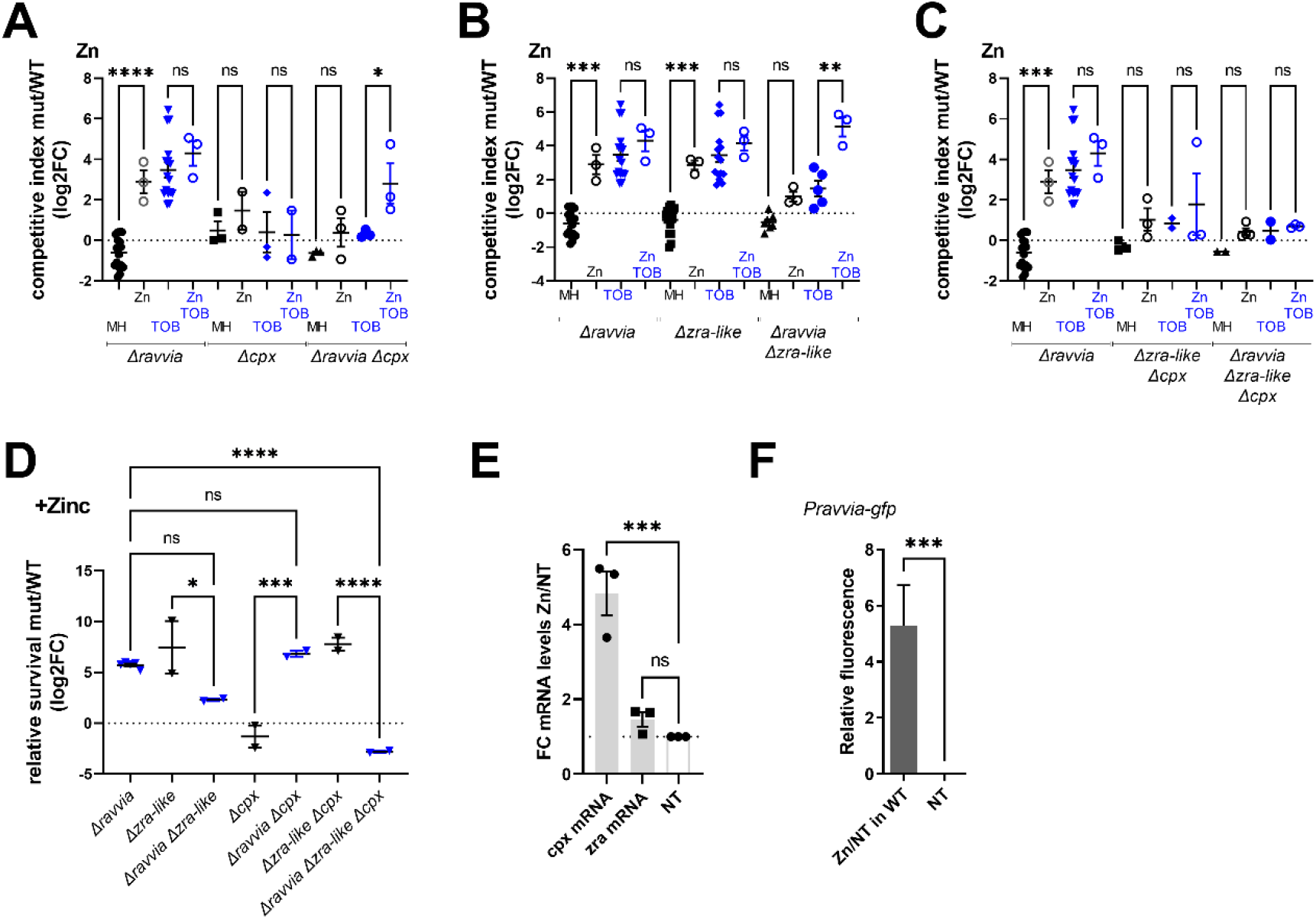
Effect of zinc supplementation. ABC. Competitions. Cultures were grown to exponential phase in MH medium supplemented with zinc during growth. “Zn” stands for ZnCl_2_: 1.5mM. The effect of deletion of *cpx* (**A**), or *zra-like* (**B**), or both (**C**) on competitive index in MH without and with TOB, where MH is the untreated growth medium. *In vitro* competition experiments of *V. cholerae* WT and indicated mutants in specified media: in black: MH: no antibiotic treatment. In blue: TOB: tobramycin 0,6 µg/ml. The Y-axis represents log_2_ of competitive index value calculated as described in the methods. A competitive index of 1 (i.e. log2 value of 0) indicates equal growth of both strains**. D. Survival** of WT and *Δravvia* to 3-hours treatment with lethal TOB at 5x MIC 5 µg/ml, in presence of zinc. The Y-axis represents log_2_ value of survival rates ratios, calculated as survival of the mutant over survival of the WT. A relative survival ratio of 1 (i.e. log2 = 0) indicates equal survival as the WT strain. **E. Expression of *cpx* and *zra*.** mRNA levels were measured using digital RT-PCR as explained in materials and methods. The Y-axis represents the fold change of induction in presence of zinc divided by the expression in the absence of zinc. **F. Expression from promoter of *ravvia*** was measured using fluorescent transcriptional fusion of *gfp* expressed from *ravvia* promoter, and quantified using flow cytometry. The Y-axis represents the relative fluorescence in Zn: zinc 1.5 mM over NT: non-treated. WT: wild type strain. For statistical significance calculations, we used one-way ANOVA. **** means p<0.0001, *** means p<0.001, ** means p<0.01, * means p<0.05. ns: non-significant. Number of replicates for each experiment: 3<n<6.

### *Δravvia* has a low ROS phenotype which depends on Cpx/Zra

We have previously shown that sub-MIC TOB leads to reactive oxygen species (ROS) formation in *V. cholerae*, which induces the bacterial SOS stress response [32]. However *Δravvia* is not more resistant to H2O2 (not shown). Considering that *Δravvia* is more tolerant to TOB and that *Δravvia* fails to induce SOS response in presence of TOB [15], we hypothesized that these two observations could be explained by a diminished ROS formation in *Δravvia*. We used the CellROX dye that, upon increased levels of ROS (O2 ^-^ and •OH), emits green fluorescence. We observed that lack of *ravvia* in *V. cholerae* leads to decreased ROS generation, both in the absence and presence of sub-MIC TOB (**Figure 5, MH and sub-MIC TOB).** We similarly observed that *Δravvia* produces decreased levels of ROS upon treatment with lethal doses of TOB **(Figure S4**).

**Figure 5:**
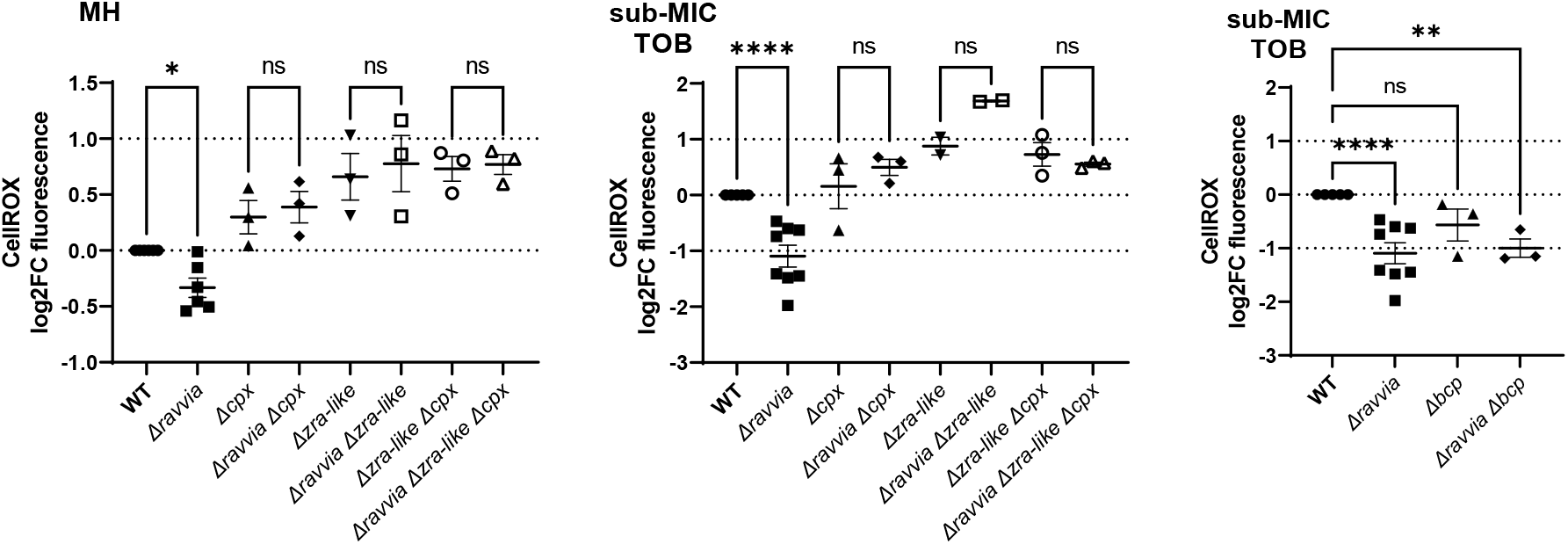
Low ROS phenotype of *Δravvia* is dependent on the presence of Cpx and Zra-like stress response systems. Quantification of variation of reactive oxygen species using CellRox. The y-axis represents log2 fold-change of detected ROS fluorescence in the indicated strain over the WT strain. Each experiment was performed at least 3 times and data and statistical significance are shown in the histograms. For statistical significance calculations, we used one-way ANOVA. **** means p<0.0001, ** means p<0.01. ns means non-significant.

We next tested whether the function of Cpx/Zra-like systems in *Δravvia* could be involved in the “low ROS” phenotype observed for *Δravvia*. **Figure 5** shows that the deletions of either *cpx* or *zra-like* (or both) suppress this phenotype, meaning that both Zra-like and Cpx are involved in the low ROS phenotype of *Δravvia*. This is consistent with the fact that both systems are also necessary for AG tolerant phenotype of *Δravvia*. As a control, we also tested the double mutant *Δravvia Δbcp*. Bcp is a thiol peroxidase responding to oxidative stress. We see no effect of *bcp* deletion on the low ROS phenotype of *Δravvia*. In conclusion, *Δravvia* shows a low ROS phenotype, which is reversed upon inactivation of Cpx and Zra-like stress responses.

### Simulatenous deletion of Cpx or Zra-like systems with *Δravvia* leads to outer membrane permeability

Since Cpx and Zra systems, have been known for their involvement in envelope stress response, we decided to test the effect of *ravvia* deletion mutant and derivatives on outer membrane permeability.

Vancomycin is an antibiotic targeting the synthesis of the peptidoglycan, but which cannot be used to treat gram-negative bacteria, because its large molecular weight prevents it from crossing the outer membrane (OM) through porins, and penetrate into the cell [33]. When the OM is damaged however, vancomycin uptake by gram negative bacteria is possible [34] [35]. We tested whether deletion of *ravvia* has an impact on vancomycin entry, by measuring the MICs of the different mutants. Our results show that single deletions of *Δravvia*, *Δcpx* or *Δzra-like* do not affect the MIC to vancomycin, while double deletions of *Δravvia* together with *Δcpx* or *Δzra-like* or both decreases the MIC from >256 µg/ml to about 64-100 µg/ml (**Figure 6A**), suggesting that simultaneous inactivation of *ravvia* function together with envelope stress response systems leads to OM damage or permeabilization.

**Figure 6:**
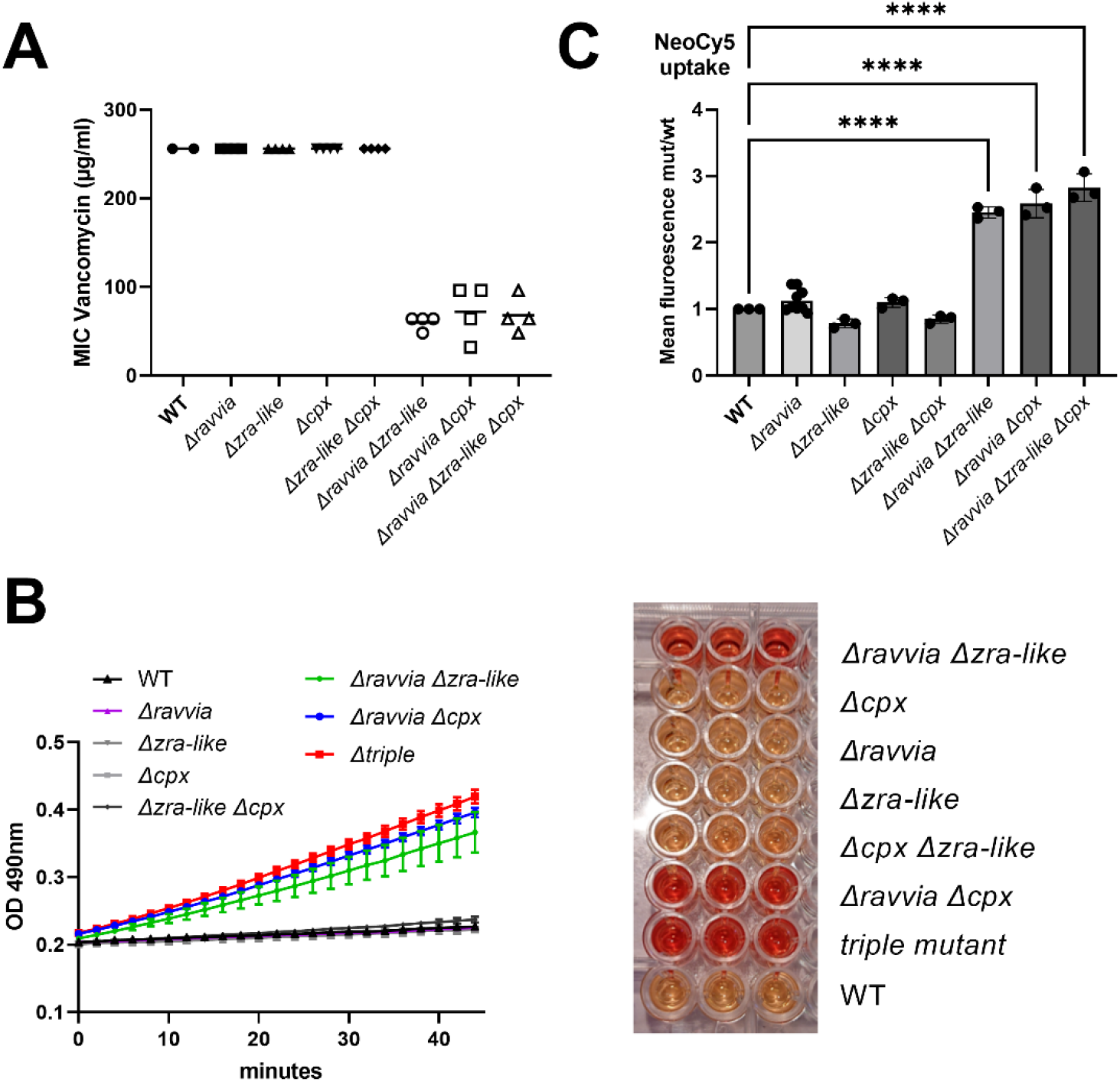
Response of *Δravvia* to the membrane targeting antibiotics. **A.** Minimum inhibitory concentration of vancomycin. MIC values are represented in the Y-axis for the indicated strains. **B.** Outer membrane permeability to nitrocefin measured on stationary phase cultures diluted to 5 × 10^7^ cells/ml by measuring the OD at 490 nm for 40 min. **C.** AG uptake quantified through Neo-cy5 entry into exponential phase cultures. Intracellular level of neomycin coupled to the fluorophore Cy5 measured by fluorescence associated flow cytometry. Error bars represent standard deviation. For statistical significance calculations, we used one-way ANOVA. **** means p<0.0001. Only significant p values are shown.

Changes in OM permeability can be quantified using nitrocefin [36], a chromogenic probe which develops color (at 490 nm) upon entry into the periplasm, in the presence of β-lactamase. We thus measured permeability of *Δravvia* and *Δcpx/Δzra-like* derivatives transformed with the low-copy pSC101 plasmid carrying the *bla* gene. Again, results show that neither single deletions of *Δravvia*, *Δcpx* or *Δzra-like,* nor the double *Δcpx Δzra-like* deletion affect nitrocefin entry, while double deletions of *Δravvia* together with *Δcpx* or *Δzra-like* or both, strongly increases it, consistent with increased OM permeability (**Figure 6B**). Since AGs also are also expected to cross the cell envelope with higher efficacy with increased OM permeability, we checked whether AG uptake is increased in *Δravvia Δcpx* or *Δravvia Δzra-like* or the triple mutant (**Figure 6C**). Consistent with vancomycin and nitrocefin entry results, neo-cy5 uptake is also increased in *Δravvia Δcpx, Δravvia Δzra-like,* but not in single mutants or the *Δcpx Δzra-like* mutant. Altogether, these results point to increased outer membrane permeability when either Cpx or Zra-like system is inactivated in *Δravvia*.

## Discussion

This study presents the first evidence of a link between RavA-ViaA function and the response to envelope stress. In fact, we identify the Cpx and a Zra-like two-component systems to be involved in the increased tolerance of *V. cholerae Δravvia* to AGs. Since Cpx and Zra systems are known to respond to protein misfolding at the periplasm, one can speculated that inactivationb of *ravvia* could generate an increase in misfolded periplasmic proteins.

In *E. coli, Δcpx* mutants show alterations in conjugational plasmid transfer, transport, ability to grow on some carbon sources and resistance to AGs. More generally, the *E. coli* Cpx transcriptome impacts inner membrane associated processes such protein secretion, and other processes like iron homeostasis, translation, and interestingly, ribosome protection factors *raiA* and *rmf* [31]. Cpx also negatively regulates respiration, energy, and TCA cycle genes. In *V. cholerae*, Cpx senses and responds to low iron, and induces iron transporters and efflux pumps [37]. Disruption of the OM and accumulation of misfolded proteins in the periplasm were shown to induce the Cpx regulon [38-40]. The fact that our TNseq data also identified the *tat* operon as important for the growth advantage of *Δravvia* in TOB also supports the existence of a so far unknown link between RavA-ViaA and misfolded proteins in the periplasm. Cpx activity is regulated by successive phosphorylations (except for *cpxP*): envelope stress leads to phosphorylation and activation of CpxA, which phosphorylates and activates CpxR [41]. Envelope stress in parallel inactivates the repressor CpxP, because CpxP binds misfolded membrane proteins and is thus titrated away from CpxA. Thus, the Cpx activity is not solely regulated by activation of the promoter.

Such protein misfolding at the periplasm can happen during AG treatment, and Cpx was in fact previously linked to AG resistance [7, 42]. Interestingly, Cpx was also shown to induce the heat-shock sigma factor RpoH in the presence of the AG gentamicin, indicating that the effect AG gentamicin at the membrane is sufficient to trigger the response to misfolded IM protein stress by Cpx [43]. *ΔcpxR* is more susceptible to AGs in *E. coli* [44], and in *Salmonella* independently of efflux pumps [45], and also independently of oxygen consumption or PMF [44], but involves altered protein composition at the membrane [46]. The AG gentamicin was shown to activate the Cpx response [47]. As a corollary, activation of Cpx leads to increased AG resistance. This was shown to be due to protection against AGs at the membrane [31, 48], partly through regulation of protein degradation at the inner membrane [46] and partly to downregulation of electron transport chains and iron import [31]. Cpx was in fact described to repress *nuo* and *cyo* aerobic respiratory complexes in *E. coli* [44], which could be toxic in conditions challenging membrane integrity, and such repression allows bacterial cells to adapt to conditions disrupting membrane integrity, among which AG induced protein misfolding. There are no described sequence homologues for *nuo* or *cyo* operons in *V. cholerae*. The functional equivalent may be the *nqr* system [49](VC2291-2-3-4), for which we see that inactivation is beneficial in WT TOB but not in *Δravvia* TOB, maybe because it is already down-regulated (approximately 2-fold but with *p-value*>0,05) in *Δravvia*.

Since the Cpx response is involved in the biogenesis of large complexes present at the envelope (respiratory complexes, but also type 4 pili, and maybe others), one could envisage that the action of Cpx may be through reduced protein trafficking at the inner membrane, hence reduced membrane stress. These properties - decrease of respiration and iron import by Cpx - can explain how ROS formation increases in *Δravvia* upon *cpx* deletion, because ROS are mainly produced with oxygen and iron through the Fenton reaction. Cpx is also closely linked to energy status of the cell and regulates protein folding and degrading factors, which are involved in adaptation to stress caused by high level of respiration[50].

In *E. coli*, ZraPSR is involved in resistance to several drugs and is important for membrane integrity[24]. Zra chaperone activity is enhanced by zinc. There is often significant overlap between processes affected by Zinc and by ROS, with zinc having mostly antioxidant function [51]. *Δzra* has increased membrane disruption during treatment with membrane targeting antibiotics. ChIP-seq and transcriptomic studies found that Zra controls the expression of *acr*, *raiA*, *rpoH*, etc. In *V. cholerae*, we find that the operon VC_1314-VC1315-VC1316 is highly upregulated in *Δravvia.* Although VC_1314 (487 aa) does not present any sequence homology to neither *zra* nor *cpx* systems, VC_1315 (449 aa) and VC_1316 (158 aa) present homologies respectively with *E. coli zraS* (465 aa) and *cpxR* (232 aa). Since the *V. cholerae* VC_1314-VC1315-VC1316 *zra*-like system also responds to zinc, we propose to name it *zraP2-zraS2-zraR2*.

One question remains open: why does deletion of the RavA-ViaA function activate the Cpx/Zra2 systems? The answer could come from the fact that when both RavA-ViaA and Cpx or Zra2 are inactivated, membrane permeability increases as observed with vancomycin and nitrocefin entry. The mechanism could be a complex one, since we also observe that RNA modifications, which impact translation [18], also impact *Δravvia* phenotypes. Note that *V. cholerae* harbors two *groESL* operons, with *gro1* being essential and *gro2* accessory. We have recently shown the involvement of *gro2* in the response to AGs [52]. Here, even in the absence of AGs, *gro2* is already essential for *Δravvia*. A specific target of this chaperone may be important in the absence of *ravvia*. *Δravvia* increases resistance to AGs but the importance of *gro2* and *clpS* may point to the existence of endogenous membrane protein stress in Δ*ravvia*. Thus, the presence of RavA-ViaA seems to be useful in order to maintain envelope integrity, and its function as a barrier against the entry of exogenous agents, such as the last resort antibiotic vancomycin.

## Acknowledgements.

We thank Louna Fruchard for her help with flow cytometry experiments. We thank Frédéric Barras for critical reading of the manuscript, Jessica El Khoury and Eduardo Rocha and Béatrice Py for valuable discussions, and Dominique Fourmy for the gift of Neo-Cy5. We also thank, for RNA-seq experiments, E. Turc, L. Lemée, T. Cokelaer, Biomics Platform and C2RT (Institut Pasteur, Paris, France, supported by France Génomique (ANR-10-INBS-09) and IBISA. We thank Sebastian Aguilar Pierlé for help with TN-seq analysis.

This research was funded by the Institut Pasteur, the Centre National de la Recherche Scientifique (CNRS-UMR 3525), the Institut Pasteur grant PTR 245-19, ANR ModRNAntibio (ANR-21-CE35-0012), ANR-LabEx [ANR-10-LABX-62-IBEID], the Fondation pour la Recherche Médicale (FRM EQU202103012569). AB was funded by Institut Pasteur Roux-Cantarini fellowship.

## Materials and methods

Table S1 shows strains used in this study and their construction.

Table S2 shows primer sequences.

### Media and Growth Conditions

Platings were done at 37°C, in Mueller-Hinton (MH) agar media. Liquid cultures were grown at 37°C in MH in aerobic conditions, with 180 rotations per minute.

**Competition experiments** were performed as described[18]: overnight cultures from single colonies of mutant *lacZ*- and WT *lacZ*+ strains were mixed 1:1 (500 μl + 500 μl). At this point 100 μl of the mix were serial diluted and plated on MH agar supplemented with X-gal (5-bromo-4-chloro-3-indolyl-β-D-galactopyranoside) at 40 μg/ml to assess T0 initial 1:1 ratio. At the same time, 5 μl from the mix were added to 200 µl of MH or MH supplemented with sub-MIC antibiotics (concentrations, unless indicated otherwise: TOB: tobramycin 0.6 μg/ml; GEN: 0.5 µg/ml; CIP: ciprofloxacin 0.01 μg/ml, CRB: carbenicillin 2.5 μg/ml), PQ: paraquat 10 µM, or H2O2: 2 mM. Cultures were incubated in 96 well plates with agitation at 37°C for 24 hours, and then diluted and plated on MH agar plates supplemented with X-gal. Plates were incubated overnight at 37°C and the number of blue and white CFUs was assessed. Competitive index was calculated by dividing the number of white CFUs (*lacZ*-strain) by the number of blue CFUs (*lacZ*+ strain) and normalizing this ratio to the T0 initial ratio.

### MIC determination

Stationary phase cultures grown in MH were diluted 20 times in PBS, and 300 μl were plated on MH plates and dried for 10 minutes. Etest straps (Biomérieux) were placed on the plates and incubated overnight at 37°C.

**Survival/tolerance tests** were performed on early exponential phase cultures. The overnight stationary phase cultures were diluted 1000X and grown until OD 600 nm of 0.35 to 0.4, at 37°C with shaking, in Erlenmeyers containing 25 ml fresh MH medium. Appropriate dilutions were plated on MH plates to determine the total number of CFUs in time zero untreated cultures. 5 ml of cultures were collected into 50 ml Falcon tubes and treated with lethal doses of desired antibiotics (5 or 10 times the MIC: tobramycin 5 or 10 μg/ml, carbenicillin 50 μg/ml, ciprofloxacin 0.025 μg/ml) for 30 min, 1 hour, 2 hours and 4 hours if needed, at 37°C with shaking in order to guarantee oxygenation. Appropriate dilutions were then plated on MH agar without antibiotics and proportion of growing CFUs were calculated by doing a ratio with total CFUs at time zero. Experiments were performed 3 to 8 times.

**Quantification of fluorescent neomycin uptake** was performed as described[28]. Neo-Cy5 is an aminoglycoside coupled to the fluorophore Cy5, and has been shown to be active against Gram-bacteria[27, 53]. Briefly, overnight cultures were diluted 100-fold in rich MOPS (Teknova EZ rich defined medium). When the bacterial strains reached an OD 600 nm of ∼0.25, they were incubated with 0.4 μM of Cy5 labeled neomycin for 15 minutes at 37°C. 10 μl of the incubated culture were then used for flow cytometry, diluting them in 250 μl of PBS before reading fluorescence. WT *V. cholerae*, was incubated simultaneously without Neo-Cy5 as a negative control. Flow cytometry experiments were performed as described[54] and repeated at least 3 times. For each experiment, 100,000 events were counted on the Miltenyi MACSquant device.

### PMF measurements

Quantification of PMF was performed using the Mitotracker Red CMXRos dye (Invitrogen) as described[29], in parallel with the neo-Cy5 uptake assay, using the same bacterial cultures. 50 µl of each culture were mixed with 60 µl of PBS. Tetrachlorosalicylanilide TCS (Thermofischer), a protonophore, was used as a negative control with a 500 µM treatment applied for 10 minutes at room temperature. Then, 25 nM of Mitotracker Red were added to each sample and let at room temperature for 15 minutes under aluminium foil. 20 μL of the treated culture were then used for flow cytometry, diluted in 200 μL of PBS before reading fluorescence.

### ROS measurements

Overnight cultures were diluted 1000X in MH medium and grown until an OD 600 nm of 0.3. Then, 100 µl of each culture was transferred to a 96-well plate, and treated with 1 µl of 250 µM CellRox Green (Thermofischer Scientific), for 30 minutes at 37 degrees, under aluminium foil. For flow cytometry, 10 µl were mixed into 200 µl of PBS. Fluorescence per cell was read on 100000 events, on the MACSquant device at 488 nm.

### RNA purification and RNA-seq

Cultures were diluted 1000X and grown in triplicate in MH to an OD 600 nm of 0.4. Frist, 1.5 ml of TriZol reagent was added to 500 µl of culture pellet followed by the addition of 300 µl of chloroforme. After centrifugation, the upper phase was mixed with a 1:1 volume of 70% ethanol before column purification. RNA was purified with the RNAeasy mini kit (Qiagen) according to manufacturer instruction (from step 4 of the protocole Part 1). Quality of RNA was controlled using the Bioanalyzer. Sample collection, total RNA extraction, library preparation, sequencing and analysis were performed as previously described [55].

### Transposon insertion sequencing

**L**ibraries were prepared as previously described [15, 56]. to achieve a library size of 600.000 clones, and subjected to passaging in MH and MH+TOB 0.5 or MH+CIP 0,001 for 16 generations [19]. A saturated mariner mutant library was generated by conjugation of plasmid pSC1819 from *E .coli* to *V. cholerae* WT. Briefly, pSC189 [15, 56] was delivered from *E. coli* strain 7257 (β2163 pSC189::spec, laboratory collection) into the *V. cholerae* WT strain. Conjugation was performed for 2 h on 0.45 µM filters. The filter was resuspended in 2 ml of MH broth. Petri dishes containing 100 µg/ml spectinomycin were then spread. The colonies were scraped and resuspended in 2 ml of MH. When sufficient single mutants were obtained (>600,000 for 6X coverage of non-essential regions), a portion of the library was used for gDNA extraction using Qiagen DNeasy Blood & Tissue Kit as per manufacturer’s instructions. This was used for library validation through insert amplification by nested PCR using a degenerate primer (ARB6), which contains 20 defined nucleotides followed by a randomized sequence. This was combined with a primer anchored in the edge of the transposon sequence (MV288) [15, 19]. After this, primer ARB3, which contains the first 20 nucleotides of ARB6 was used for nested amplification in combination with MV288. After validation, the libraries were passaged in MH media for 16 generations with or without 50%MIC of TOB or CIP, in triplicate. gDNA from time point 0 and both conditions after 16 generation passage in triplicate was extracted.

Sequencing libraries were prepared using Agilent’s SureSelect XT2 Kit with custom RNA baits designed to hybridize the edges of the Mariner transposon. The 100 ng protocol was followed as per manufacturer’s instructions. A total of 12 cycles were used for library amplification. Agilent’s 2100 Bioanalzyer was used to verify the size of the pooled libraries and their concentration. HiSeq Paired-end Illumina sequencing technology was used producing 2×125 bp long reads. Reads were then filtered through transposon mapping to ensure the presence of an informative transposon/genome junction using a previously described mapping algorithm [57]. Informative reads were extracted and mapped. Reads were counted when the junction was reported as mapped inside the CDS of a gene plus an additional 50 bp upstream and downstream. Expansion or decrease of fitness of mutants was calculated in fold changes with normalized insertion numbers. Normalization calculations were applied according to van Opijnen et al [58]. Expansion or decrease of fitness of mutants was calculated in fold changes with normalized insertion numbers. Baggerly’s test on proportions [59] was used to determine statistical significance as well as a Bonferroni correction for multiple hypotheses testing.

### mRNA quantifications by digital-RT-PCR

qRT-PCR reactions were prepared with 1 μl of diluted RNA samples using the qScript XLT 1-Step RT-qPCR ToughMix (Quanta Biosciences, Gaithersburg, MD, USA) within Sapphire chips. Digital PCR was conducted on a Naica Geode (programmed to perform the sample partitioning step into droplets, followed by the thermal cycling program suggested in the user’s manual. Image acquisition was performed using the Naica Prism3 reader. Images were then analyzed using Crystal Reader software (total droplet enumeration and droplet quality control) and the Crystal Miner software (extracted fluorescence values for each droplet). Values were normalized against expression of the housekeeping gene *gyrA* as previously described [60].

### Quantification of *gfp* fusion expression by fluorescent flow cytometry

Flow cytometry experiments were performed as described [54] on overnight cultures and repeated at least 3 times. For each experiment, 50,000 to 100,000 events were counted on the Miltenyi MACSquant device.

Transcriptional fusion: *ravvia* promoter sequence fused to *gfp* by amplification of *Pravvia-gfp* from pZE1-*gfp* [61] using primers ZIP537/ZIP200. The fragment was cloned into pTOPO-TA cloning vector. The *Pravvia-gfp* fragment was then extracted using *EcoRI* and cloned into the low copy plasmid pSC101. The plasmid was introduced into desired strains, and fluorescence was measured on indicated conditions, by counting 100,000 cells on the Miltenyi MACSquant device

### Growth on microtiter plate reader

Overnight cultures were diluted 1:500 in fresh MH medium, on 96 well plates. Each well contained 200 μl. Plates were incubated with shaking on TECAN plate reader device at 37°C, OD 600 nm was measured every 15 minutes. Tobramycin was used at sub-MIC: TOB 0.6 μg/ml. The concentrations of other antibiotics are specified on each figure.

### Quantification of nitrocefin entry

Nitrocefin, a chromogenic probe, 3-(2,4-dinitrostyryl)-(6 R,7 R)-7-(2-thienylacetamido)-ceph-3-em-4-carboxyl acid (Calbiochem), was used to assess membrane permeability. Tested strains were transformed with the low copy pSC101 plasmid carrying the *bla* gene coding for a β-lactamase, a periplasmic protein which allows coloration of nitrocefin upon entry into the bacterial cell. Cells were grown to stationary phase, washed twice and resuspended at a concentration of 5×10^7^ cells/ml in PBS. The reaction was performed with 175 μl of PBS buffer and 25 μl of the nitrocefin 0.5 mg/ml stock solution, in 96 well plates. 50 μl of the bacterial suspension was added and the OD at 490 nm was measured every 2 min for 45 min using a plate reader at 37°C, with shaking for 10 s every minute.

### Data availability

The data discussed in this publication have been deposited in NCBI’s Gene Expression Omnibus and are accessible through GEO Series accession number GSE196651 (GSM5897403, GSM5897404, GSM5897405, GSM5897412, GSM5897413, GSM5897414) for RNAseq data and GSE198341 for TNseq data (GSM5945317, GSM5945318, GSM5945319, GSM5945323, GSM5945324, GSM5945325, GSM5945329, GSM5945330, GSM5945331, GSM5945338, GSM5945339, GSM5945340, GSM5945341, GSM5945342, GSM5945343).

## Supplementary figures

**Figure S1:**
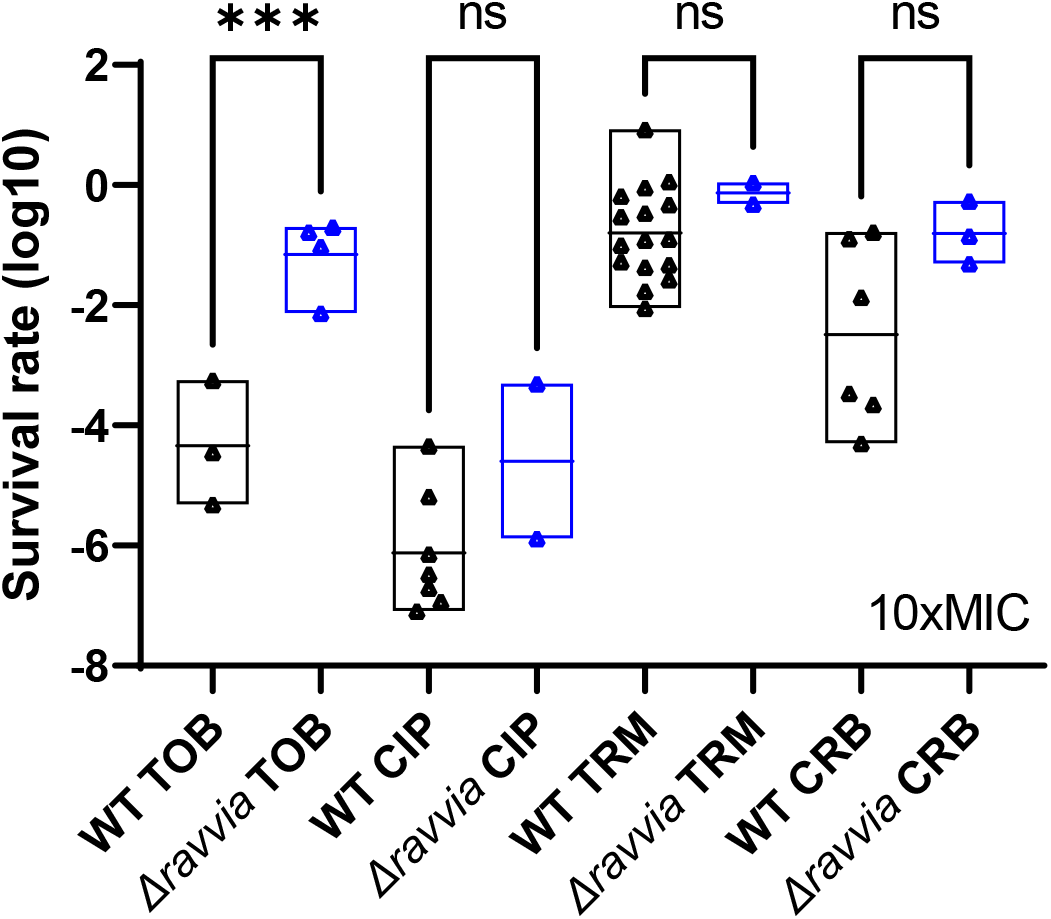
Effect of RavA-ViaA on tolerance to antibiotics. **A.** Survival of WT and *Δravvia* to treatment with antibiotics at 10x the MIC. Cultures were grown without antibiotics up to early exponential phase, and treated with antibiotics at lethal concentrations. TOB: tobramycin 10 µg/ml. Non-aminoglycoside antibiotics: CIP: ciprofloxacin. TRM: trimethoprim. CRB: carbenicillin. For statistical significance calculations, we used one-way ANOVA. *** means p<0.001, ns means non-significant. Number of replicates for each experiment: n>=3.

**Figure S2.**
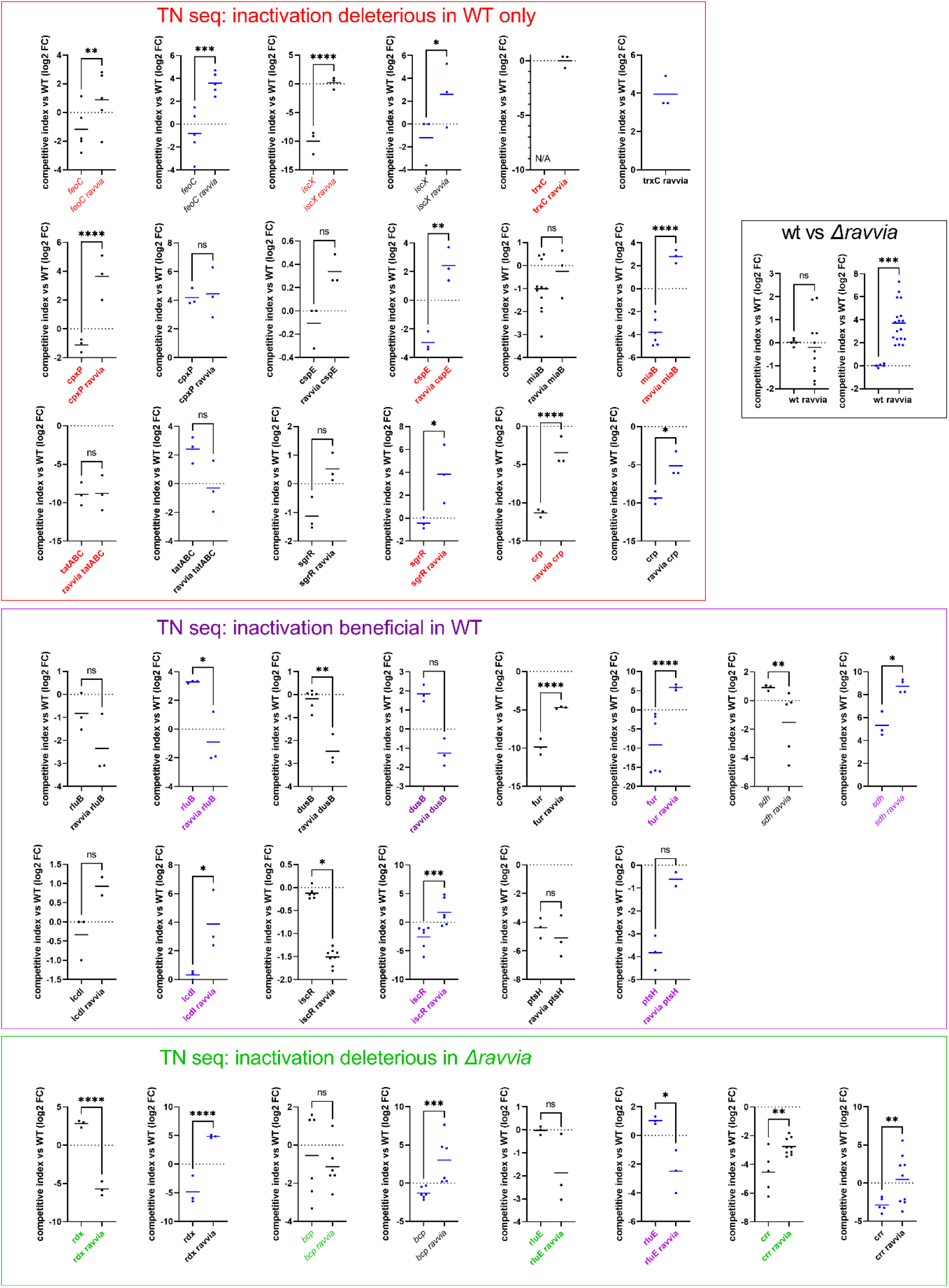
Impact of selected gene deletions in WT and *Δravvia* on fitness during growth in sub-MIC antibiotics: *in vitro* competition experiments of *V. cholerae* WT and mutant strains in the absence or presence of TOB at sub-MICs (50% of the MIC). We were looking for factors for which inactivation would lead to loss of the fitness advantage of Δ*ravvia* in AGs. We tested the effect of 21 gene deletions, selected because they are important in WT but not in Δ*ravvia* (panel with red square), important in Δ*ravvia* (green panel) or beneficial in WT only (purple panel). For 16 of them, none completely suppressed this phenotype. TN-seq screen identified functions that are necessary for AG resistance in *Δravvia*. (i) in the absence of TOB: several genes are no longer essential or as important in *Δravvia* as in WT. These genes include carbohydrate utilization and metabolism genes (*crp*), iron and respiration related factors (*feoC, iscX, trxC*), and envelope related genes (*cpxP*). This suggests that deletion of *ravvia* leads to changes in carbon metabolism, iron and respiration, and membrane stress. This is consistent with transcriptomic data. Several genes become essential or important in *Δravvia:* genes involved in respiration and iron utilization (e.g. *rdx, bcp)*; carbohydrate metabolism (*crr*), translation (*truC, rluE*) and protein stress (*groES2, clpS,* not tested here in competition). (ii) after growth with TOB: several genes are no longer needed in *Δravvia:* electron transport/redox, carbohydrate metabolism (*sgrR*), stringent response and translation stress (e.g. *raiA, hpf,* not tested here), envelope and cell division, e*.g. cspE,* coding for a cold shock transcription anti-terminator which interacts specifically with mRNAs that encode membrane proteins [32]. In summary, several functions appear to be less needed in *Δravvia*: proteins protecting ribosomes upon translation stress (e.g. hibernation factors), consistent with the that AGs cause less translation stress in *Δravvia*. For cytochromes, their inactivation probably decreases PMF and confers AG resistance in WT but since *Δravvia* is already more resistant, their effect on PMF may have little impact on AG tolerance of *Δravvia*. Notable phenotypes were conferred by deletion of the *tat* operon (export of folded proteins to the perisplasm) which leads to loss of *Δravvia’s* growth advantage in TOB, as well as RNA modification factors *dusB*, *rluB*, *rluE*, for which deletion is known to be beneficial in TOB [18], and cpxP which confers an advantage only to the WT strain. MH: no antibiotic treatment (black dots). TOB: tobramycin 0.6 μg/ml (blue dots). The Y-axis represents log_2_ of competitive index value calculated as described in the methods. A competitive index of 1 (i.e. log2 value of 0) indicates equal growth of both strains. Statistical comparisons are between the competition [*Δgene vs* WT] and [*Δgene Δravvia* vs WT]. For statistical significance calculations, we used one-way ANOVA. **** means p<0.0001, *** means p<0.001, ** means p<0.01, * means p<0.05. ns: non-significant. Number of replicates for each experiment: 3<n<6.

**Figure S3:**
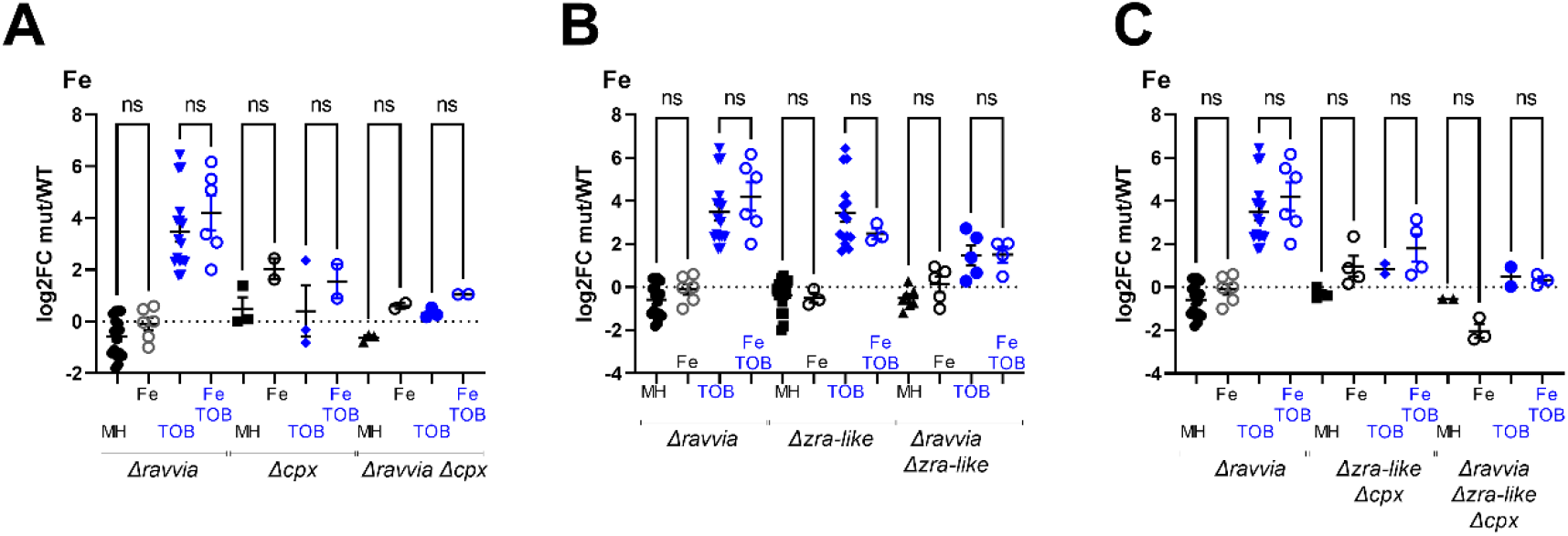
Effect of iron supplementation. ABC. Competitions. The effect of deletion of *cpx* (**A**), or *zra-like* (**B**), or both (**C**) on competitive index in MH with addition of iron, without and with TOB. Cultures were grown to exponential phase in MH medium supplemented with iron during growth. “Fe” stands for FeSO4 18µM. *In vitro* competition experiments of *V. cholerae* WT and indicated mutants in specified media: in black: MH: no antibiotic treatment. In blue: TOB: tobramycin 0,6 µg/ml. The Y-axis represents log_2_ of competitive index value calculated as described in the methods. A competitive index of 1 (i.e. log2 value of 0) indicates equal growth of both strains. For statistical significance calculations, we used one-way ANOVA. **** means p<0.0001, *** means p<0.001, ** means p<0.01, * means p<0.05. ns: non-significant. Number of replicates for each experiment: 3<n<8.

**Figure S4:**
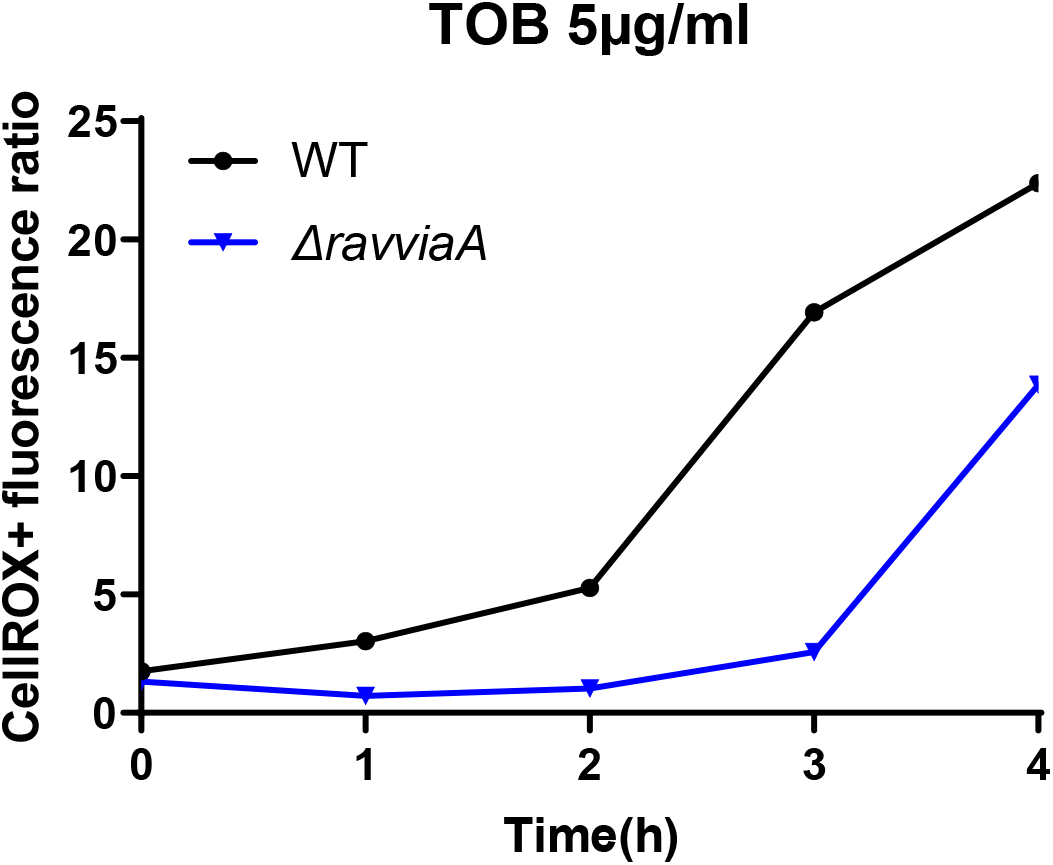
Low ROS phenotype of *Δravvia* is maintained in the presence of 5x MIC TOB. Quantification of variation of reactive oxygen species using CellRox. The y-axis represents fluorescence corresponding to detected ROS in the indicated strain, as a function of time. Experimental conditions were that of survival assays performed on exponentially growing cultures. Fluorescence was measured using flow cytometry every hour during TOB treatment, on 50,000 cells.

**Table S1:**
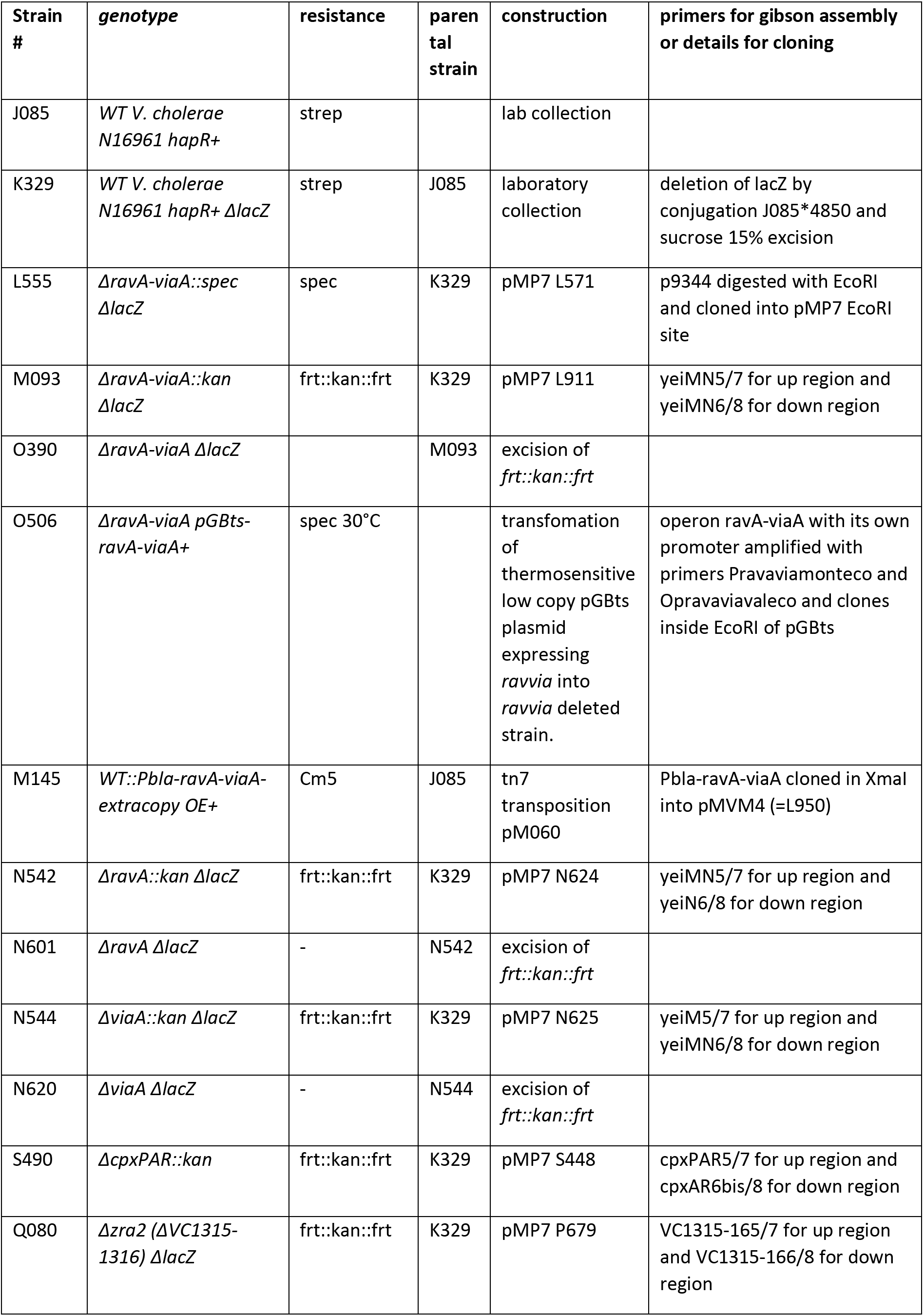
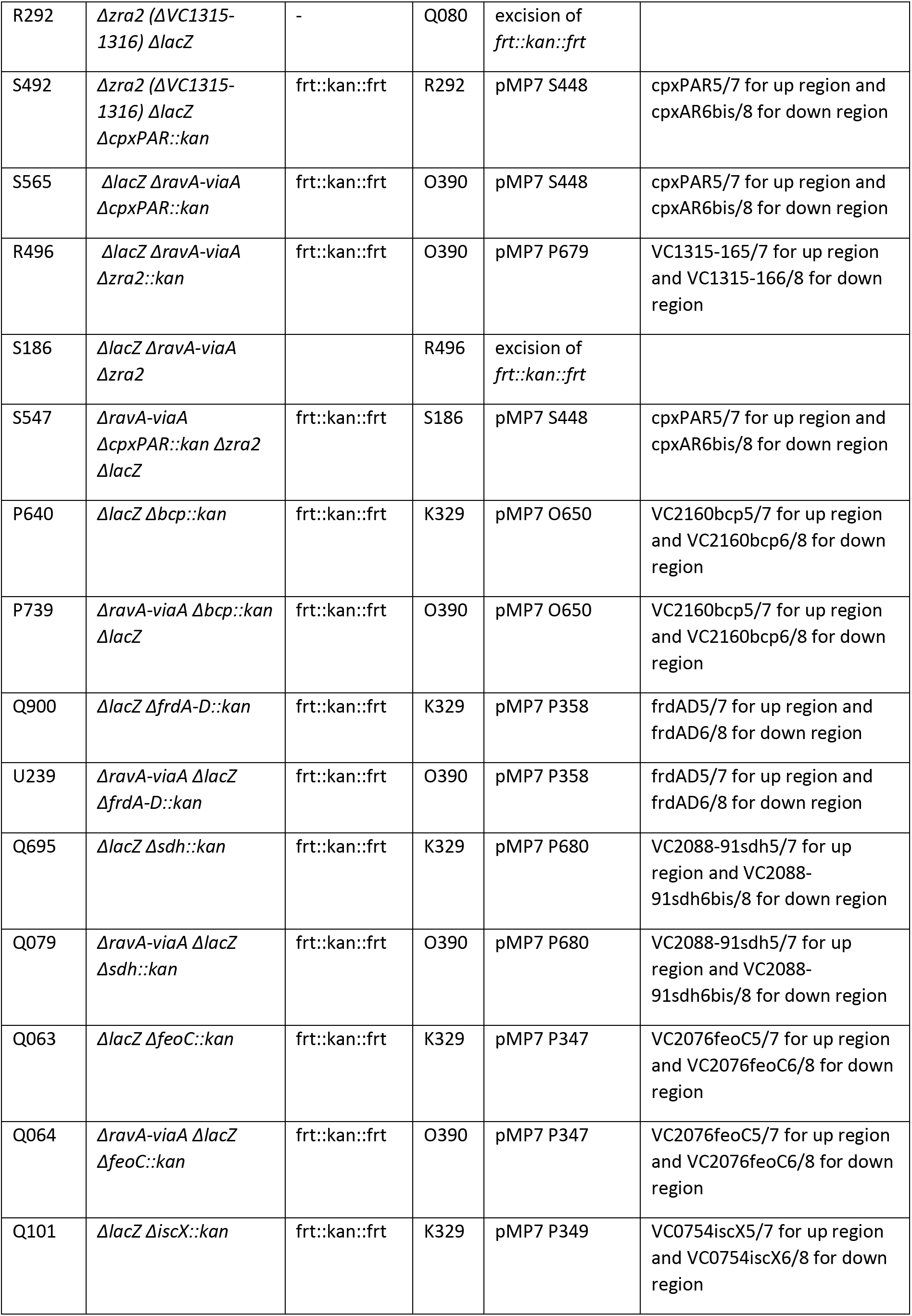
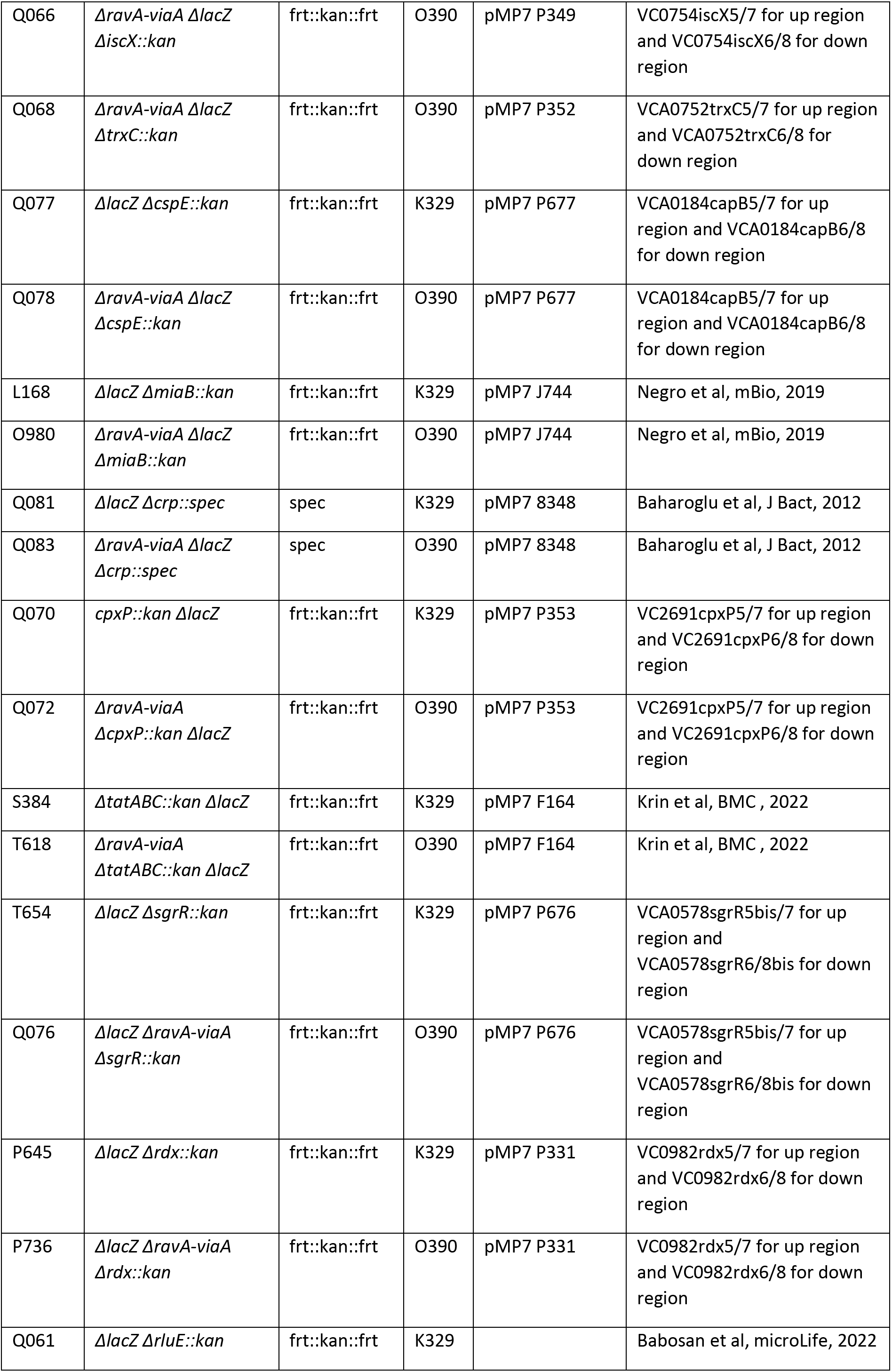
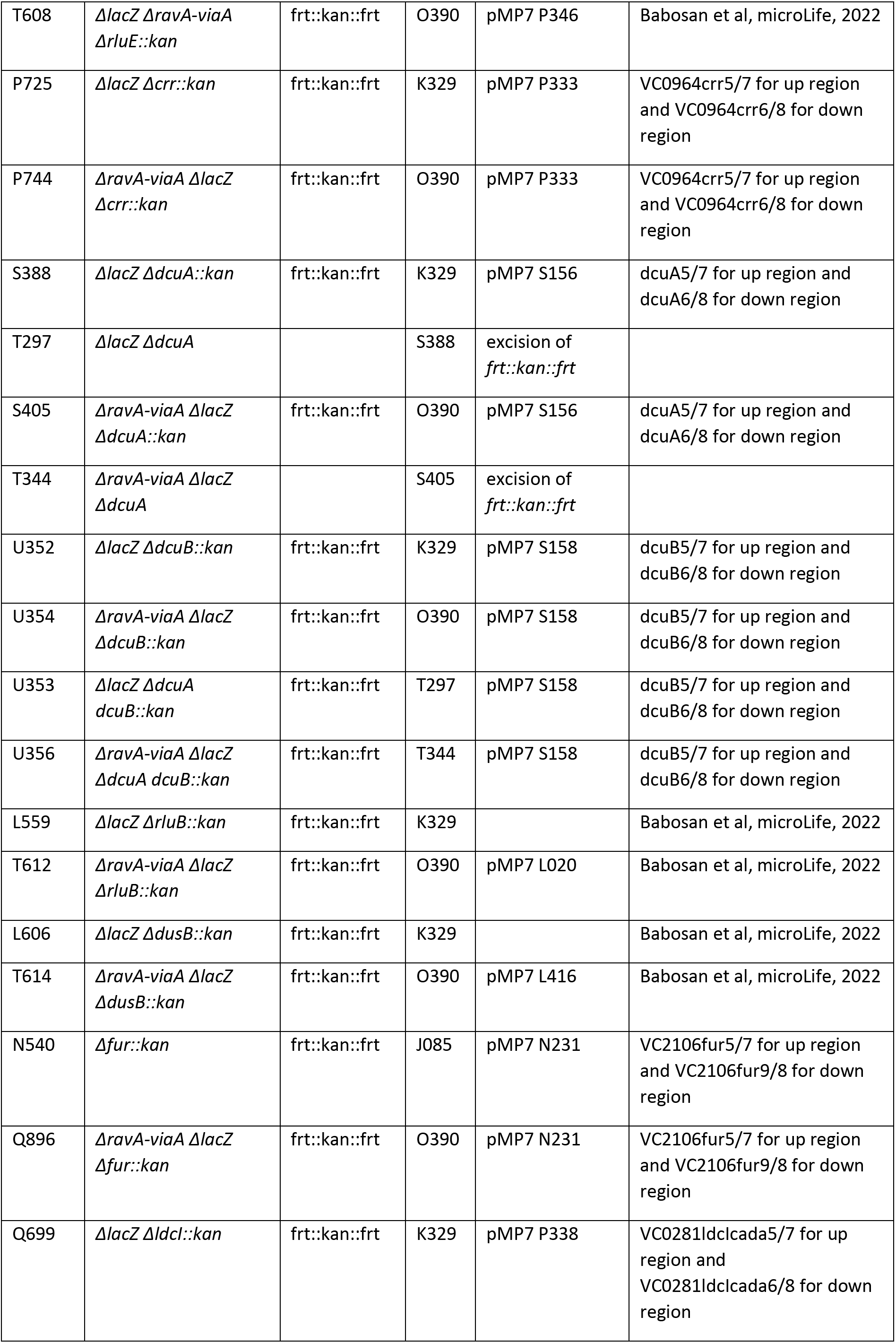
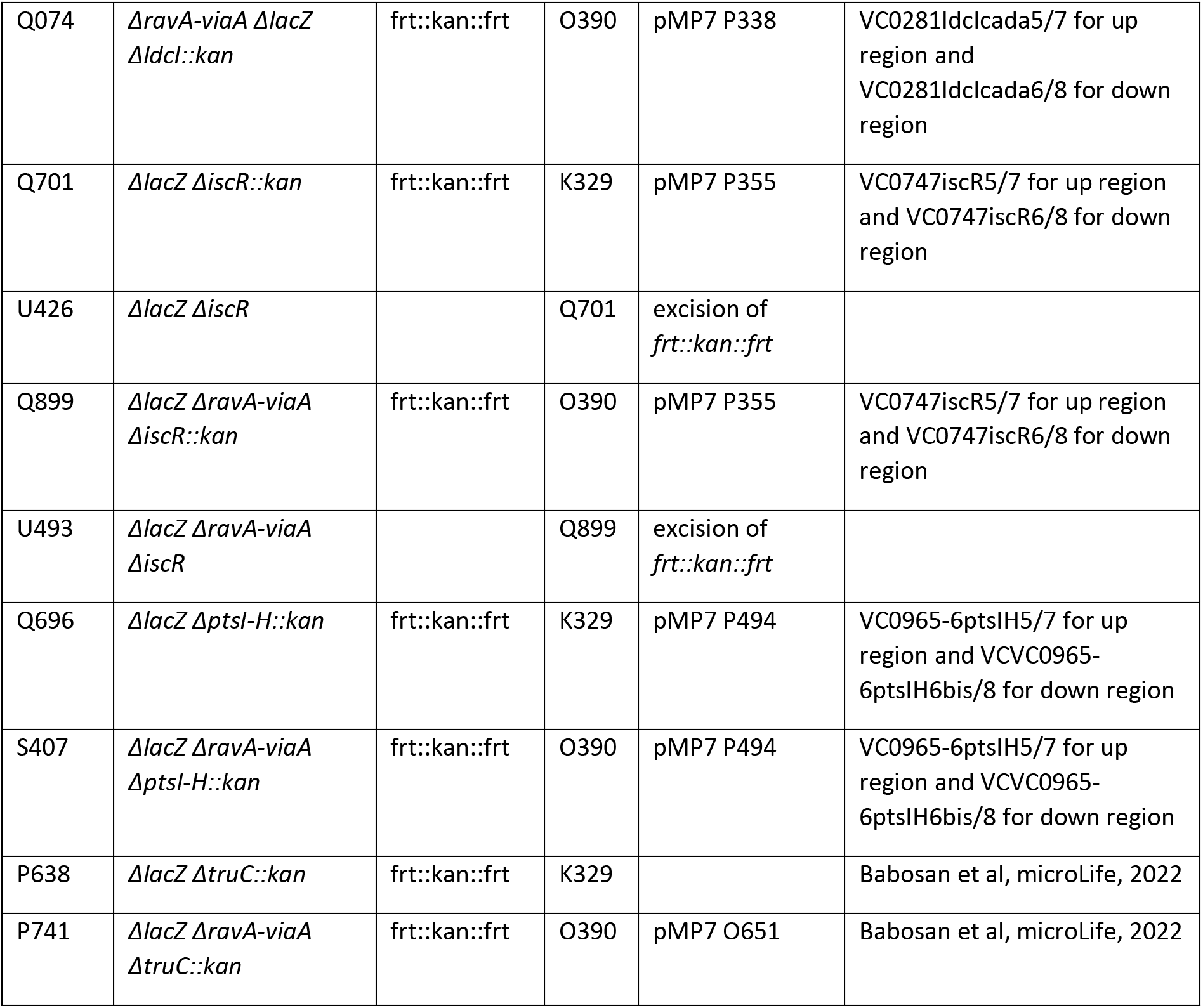
Strains.

**Table S2:**
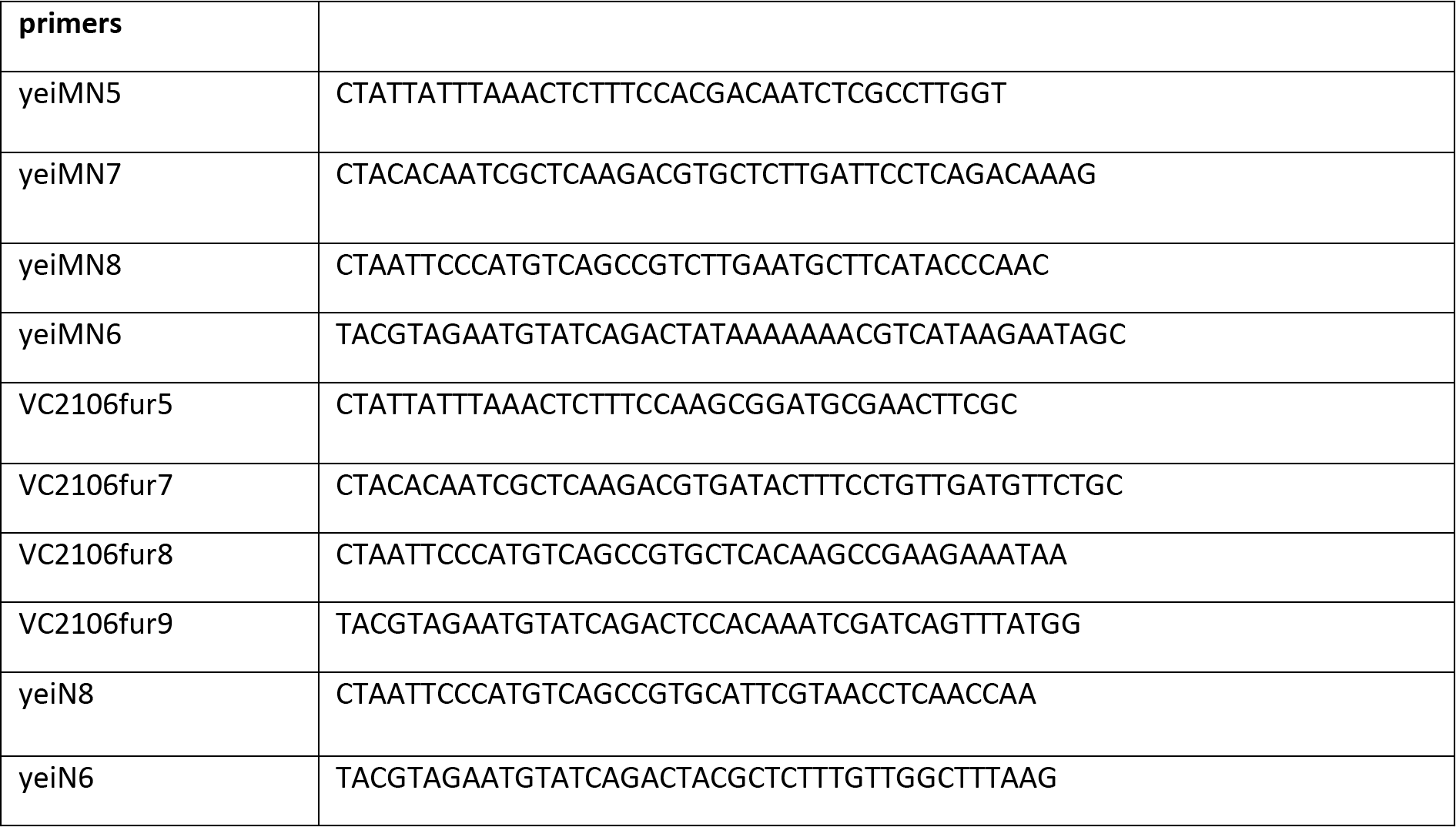
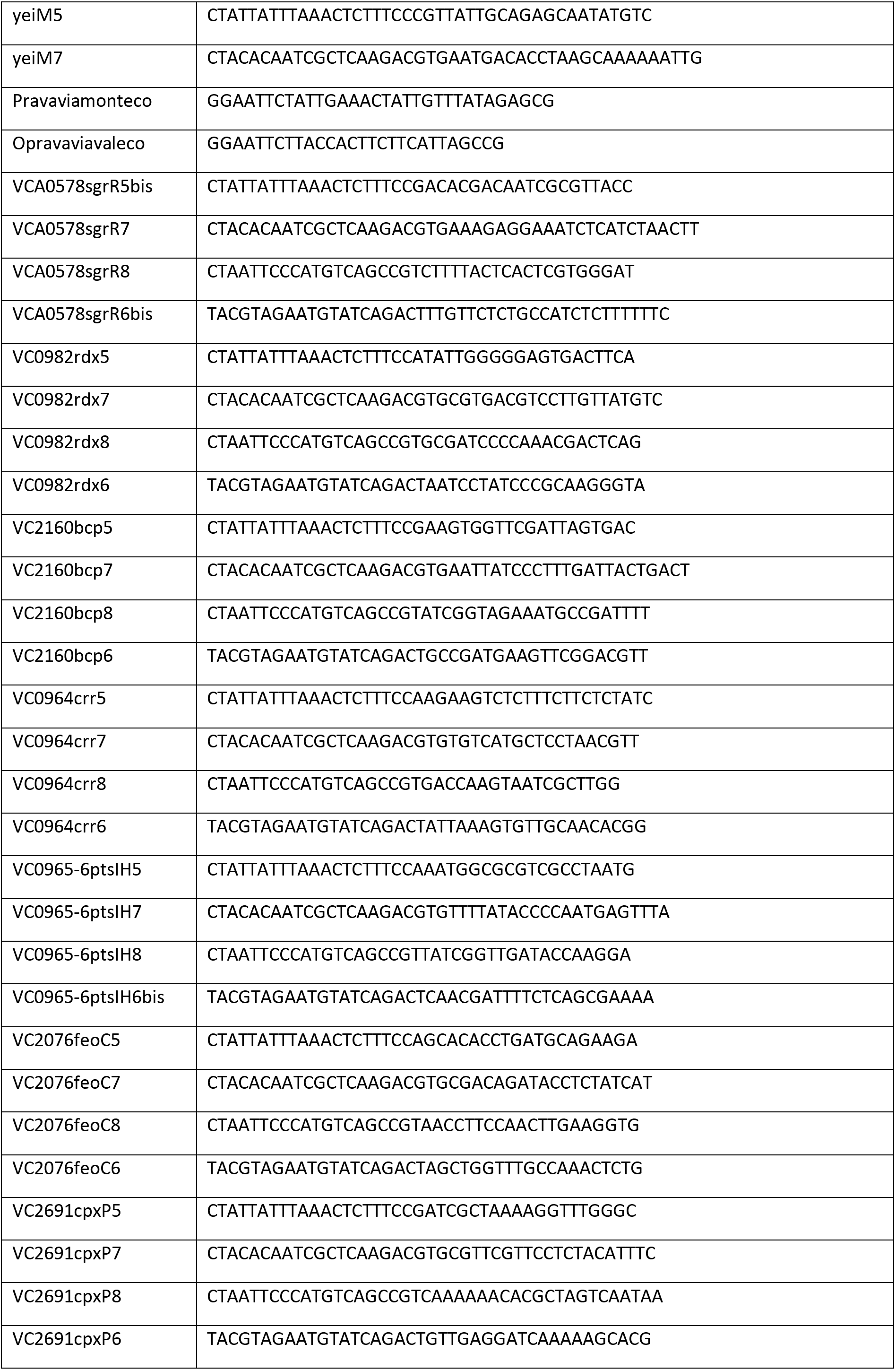
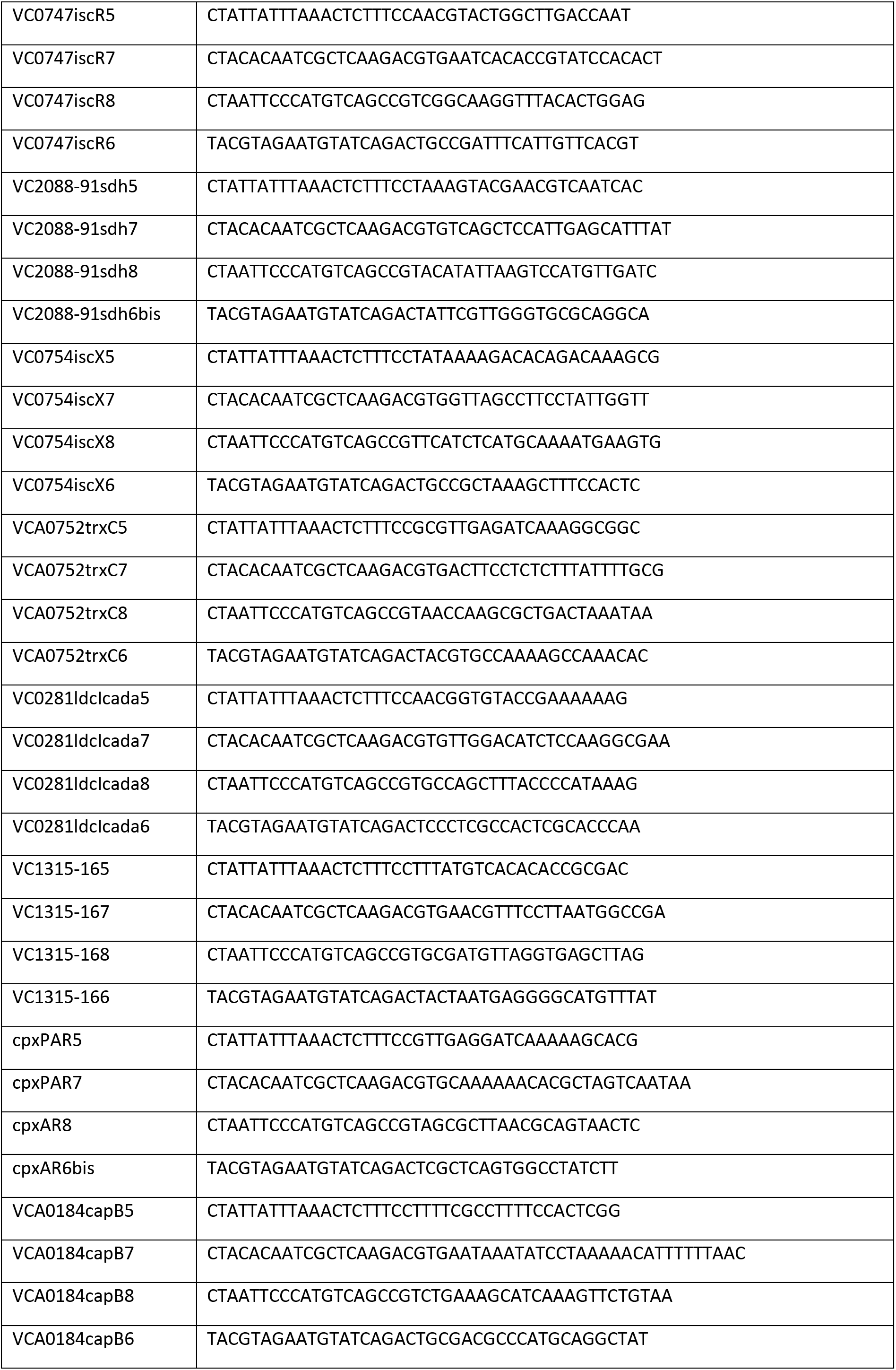
Plasmids.

